# Scrutinized lipid utilization disrupts Amphotericin-B responsiveness in clinical isolates of *Leishmania donovani*

**DOI:** 10.1101/2024.10.21.619374

**Authors:** Supratim Pradhan, Dhruba Dhar, Debolina Manna, Shubhangi Chakraborty, Arkapriya Bhattacharyya, Khushi Chauhan, Rimi Mukherjee, Abhik Sen, Krishna Pandey, Soumen Das, Budhaditya Mukherjee

**Affiliations:** School of Medical Science and Technology, Indian Institute of Technology Kharagpur, India-721302; ICMR-Rajendra Memorial Research Institute of Medical Sciences, Patna, India-800007

**Keywords:** Clinical Leishmania donovani isolates, drug-unresponsiveness, Fluid phase endocytosis /vesicle fusion/LDL-cholesterol, Lipid droplets

## Abstract

The management of *Leishmania donova*ni (LD), responsible for fatal visceral leishmaniasis (VL), faces increasing challenges due to rising drug-unresponsiveness, leading to increasing treatment failures. While hypolipidemia characterizes VL, LD, a cholesterol auxotroph, relies on host lipid scavenging for its intracellular survival. The aggressive pathology, in terms of increased organ parasite load, observed in hosts infected with antimony-unresponsive-LD (LD-R) as compared to their sensitive counterparts (LD-S), highlights LD-R’s heightened reliance on host lipids. Here we report that LD-R-infection promotes fluid-phase endocytosis in the host, selectively accumulating neutral lipids while excluding oxidized-LDL. LD-R enhances the fusion of endocytosed LDL-vesicles with its phagolysosomal membrane and inhibits cholesterol mobilization from these vesicles by suppressing NPC-1. This provides LD-R amastigotes with excess lipids, supporting their rapid proliferation and membrane synthesis. This excess LDL-influx leads to an eventual accumulation of neutral lipid droplets around LD-R amastigotes, thereby increasing their unresponsiveness towards Amphotericin-B, a second-line amphiphilic antileishmanial. Notably, VL patients showing relapse with Amphotericin-B treatment exhibited significantly lower serum LDL and cholesterol than cured cases. Treatment with Aspirin, a lipid droplet blocker, reduced lipid droplets around LD-R amastigotes, restoring Amphotericin-B responsiveness.

## Introduction

Antimony, which was mainstay for treating fatal visceral leishmaniasis (VL), has been withdrawn from Indian Subcontinent for more than a decade now due to emergence of resistance. Despite this, recent clinical LD isolates remain unresponsive to antimony (LD-R), indicating persistent phenotype [1]. Notably, LD-R also exhibits more aggressive pathology in clinical and experimental infection featuring increased organ parasite burden as compared to LD-S-infection [2–5]. Increased metacyclogenesis among LD-R was initially thought to be the primary factor contributing to higher infectivity, resulting in organ-parasite overload [6]. However, later studies revealed that LD-R promastigotes have a greater replicative potential than LD-S, possibly offering a selective intracellular survival advantage [7, 8]. This implies LD-R might be more adept at acquiring and metabolizing host derived nutrients within their parasitophorous vacuole (PV), facilitating their heightened intracellular proliferation. Lipids are crucial carbon sources for energy in proliferating intracellular pathogens. Several intracellular bacteria, protozoans and viruses utilizes cellular lipid droplets and plasma lipoproteins for entry, energy acquisition and biomembrane synthesis, demonstrating intricate interactions between intracellular pathogens and host lipid metabolism [9, 10]. *Leishmania* parasites are cholesterol auxotrophs [11], and need (i) host membrane cholesterol for successful attachment and infection [12]. (ii) Intracellular leishmania amastigotes replicating within membrane-bound PVs, accumulates cholesterol and lipid-droplets [13]. (iii) *Leishmania* amastigotes isolated from infected host have been shown to have increased cholesterol content in them [14], all these clearly suggest replicating *Leishmania* amastigotes might co-opt host derived lipids to facilitate their intracellular proliferation and membrane synthesis. In fact, hypocholesteremia is a hallmark of VL, with intracellular LD depleting host cholesterol and lipoproteins [15, 16]. However, the mechanism by which PV-bound LD amastigotes acquire this lipids, and how variations in lipid type, source, and utilization dynamics affect disease pathology and drug responsiveness, remains unclear.

So far, drug responsiveness in leishmaniasis has mostly been studied through in vitro selection of drug-resistant promastigotes, with limited focus on the drug susceptibility profiles of circulating clinical strains of *Leishmania* [17–19]. However, growing evidences indicates that drug responsiveness and treatment failure in VL are multifactorial, influenced by the host’s immune-metabolic status [3, 20]. Importantly, recent reports have also linked emergence of drug unresponsiveness among intracellular pathogens with their ability to modulate host lipid metabolism [21, 22]. For instance, increased lipid accumulation can hinder drug penetration by trapping lipophilic drugs, fostering resistance [23–25]. VL treatment in recent years have seen a rise in treatment failure against Amphotericin-B (Amp-B), a second-line lipophilic drug [2], making it critical to investigate these failures in the context of lipid utilization during LD infection.

Using clinical antimony unresponsive LD-strains isolated from patients this study elucidates how LD-R-infection leads to an increased uptake of low-density lipoprotein (LDL) leading to eventual accumulation of neutral lipids droplets within their PV. In vitro infection of liver-resident Kupffer cells (KCs), primary sites for cholesterol synthesis and LD amastigote residence, demonstrated that LD-R infection facilitates fluid-phase endocytosis, increasing LDL-vesicle uptake in infected-KCs cells. LD-R-PV exhibits a higher propensity to fuse with endocytosed LDL-vesicles, coupled with the inhibition of cholesterol recycling from the vesicles back to the plasma membrane of Infected-MFs. Our observation thus explains the heightened energy source and cholesterol availability for PV and daughter membrane synthesis to accommodate higher number of LD-R amastigotes observed in clinical infection. We also observed a suppressed expression of MSR-1, a scavenger receptor of ox-LDL [26], in LD-R infected murine liver, suggesting LD-R-infection limits the uptake of ox-LDL, reducing inflammatory responses. Furthermore, our findings indicate that lipid droplet accumulation around LD-R amastigotes contributes to Amp-B unresponsiveness, explaining recent reports of Amp-B cross-resistance in clinical LD-R isolates from India [27, 28]. Significant low lipid profiles in VL patients experiencing treatment failure against liposomal Amp-B, combined with complete clearance of LD-R amastigotes using a combination treatment of Aspirin (a lipid droplet formation blocker) and Amp-B, confirmed critical role of assimilating host lipids in developing Amp-B unresponsiveness among rapidly replicating LD amastigotes exhibiting primary unresponsiveness towards antimony. This study identifies sequence of events that enables LD-R amastigotes to redirect host nutrients for their own aggravated sustenance, and how this preferential lipid utilization might equip them with unresponsiveness against Amp-B contributing to the gradual spread of cross-resistance among clinical LD-R isolates.

## Results

### 1. LD-R infection caused severe dyslipidaemia as compared to LD-S infection

Intracellular amastigotes of *Leishmania* parasites exhibit lipid auxotrophy [11], with dyslipidaemia being a hallmark for leishmaniasis [15], including VL patients. Previously, LD amastigote load in splenic aspirates of VL patients have been shown to exhibit an inverse correlation with serum cholesterol level [29]. Since infection with antimony-unresponsive-LD (LD-R) has been previously reported to exhibit higher amastigote load in spleen and liver of infected host, as compared to infection with their responsive counterparts (LD-S) [7, 30], we wanted to investigate possible extent of dyslipidaemia in response to LD-R and LD-S-infection and how it might affect disease pathology. Mice infected with equal number of sorted metacyclics (**Fig S1A**) of natural clinical LD-isolates with confirmed LD-R phenotype, MHOM/IN/09/BHU575/0 (LD-R^1^) and MHOM/IN/10/BHU814/1 (LD-R^2^), showed a significantly lower serum levels of LDL, HDL, and cholesterol as compared to MHOM/IN/83/AG83 (LD-S^1^) and MHOM/IN/80/DD8 (LD-S^2^), infected mice (**Figure 1A).** Liver serves as a primary site for lipid synthesis and liver-resident MFs (Kupffer cells, KC) are one of the primary sites harboring LD amastigotes [31]. Therefore, we decided to test if KCs can be established as an *in vitro* model for LD-infection to investigate the possible link of host lipid utilization with respect to aggressive pathology shown by LD-R. Purified KCs (**Figure S1B**) infected with equal number of sorted metacyclics for LD-S and LD-R-isolates as before displayed no significant difference in their initial attachment to host membrane (**Figure 1B** and **Movie 1A, 1B).** However, although initial number of intracellular LD-S and LD-R-amastigotes 4Hrs post infection (p.i) showed no difference, a significantly higher number of LD-R amastigotes were observed in 24Hrs infected-KC (**Figure 1Ci, ii**). Interestingly, as compared to a significant and sharp rise in the number of intracellular amastigotes between 4Hrs and 24Hrs post infected KC in response to LD-R infection, the number of intracellular amastigotes although increased significantly did not doubled from 24Hrs to 48Hrs p.i. suggesting exponential LD amastigote replication between 4Hrs and 24Hrs time frame and slowing down after that (**Figure 1Ci, ii**). Moreover, it was also noticed that at 72Hrs p.i. a notable number of infected-KC began detaching from the wells with extracellular amastigotes probably egressing out from the infected-KCs (**Movie 2**). Thus, 24Hrs time point was selected to conduct all further infection studies involving KCs. Parallel infection performed in peritoneal exudate macrophages (PEC), which serves as a long-standing model for *in vitro* LD-infection, under identical experimental conditions also resulted in similar observation with LD-R strains showing significantly higher amastigote load at 24Hrs p.i than LD-S strains (**Figure S1Ci, ii, iii**), as previously reported by multiple groups [1, 4, 6]. Moreover, KCs infected with LD-S or LD-R amastigotes isolated from 28 days infected mouse spleens, also resulted in significantly higher parasite load for LD-R (**Figure S1D**), clearly indicating that higher parasite load is a conserved LD-R phenotype which does not depends on initial infectivity or MF origin. Notably, infected-KCs had a slightly higher inherent amastigote load compared to infected-PECs for both LD-S and LD-R-infections, with no significant difference in parasite load within the two independent LD-S or LD-R strains (**Figure 1C, Figure S1C, S1D**). Hence, BHU575 was selected as representative LD-R and AG83 as representative LD-S to conduct all further comparisons with the inclusion of other representative strains as and when mentioned. Higher intracellular amastigote load in response to LD-R-infection was further confirmed by imaging LD-S and LD-R-PV at 4Hrs and 24Hrs infected-KCs (**Figure 1D**). Comparative analysis revealed significantly higher number of LD-R-PVs as compared to LD-S-PVs at 24Hrs p.i, although this difference was not observed at 4Hrs p.i (**Figure 1Di, Dii**). Interestingly, 3D image reconstruction for LD-R-PV at 24Hrs p.i, presented it as a larger fused structure as compared to smaller individual segmented appearance of LD-S-PV (**Figure 1Diii**).

**Figure 1.**
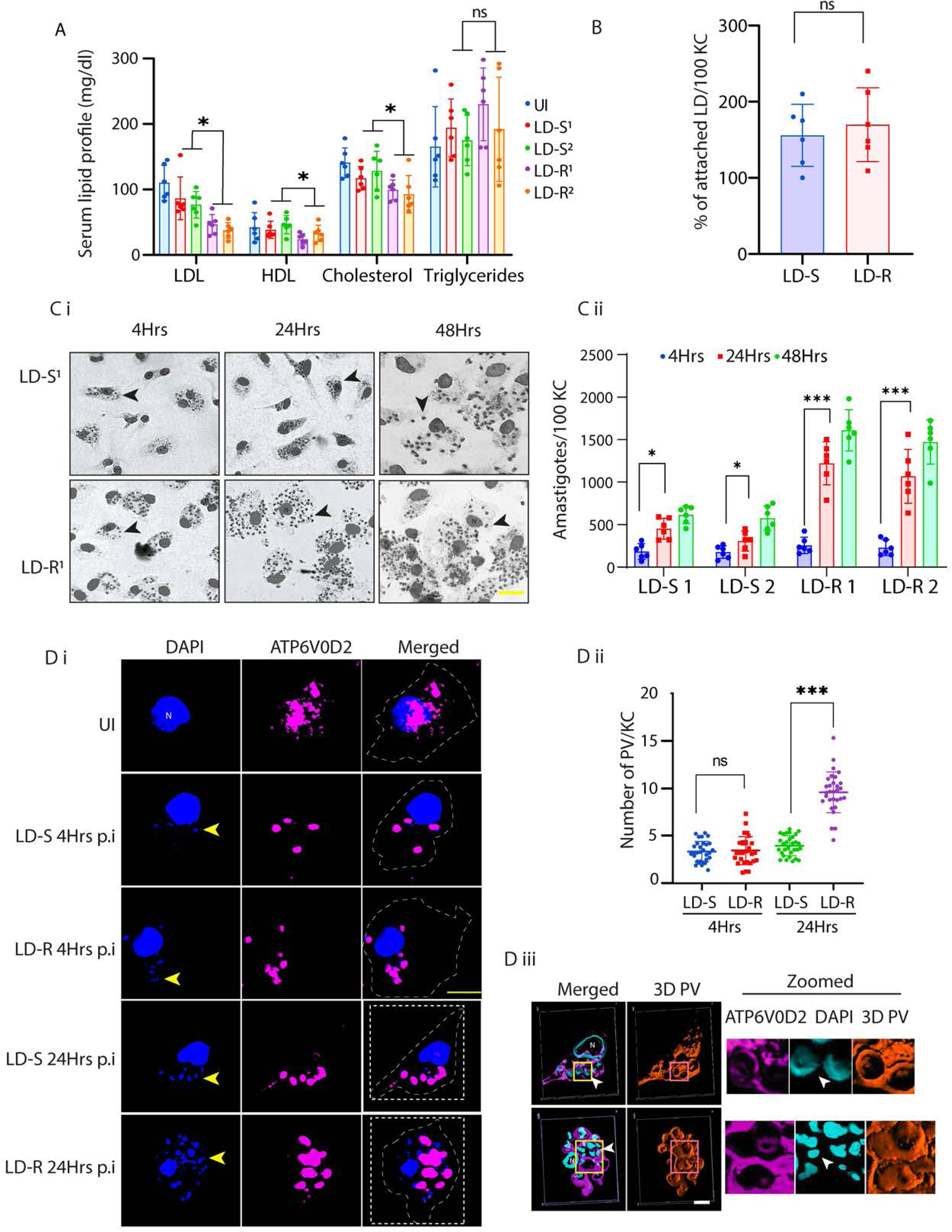
Severe dyslipidemia is linked with higher intracellular proliferation of LD-R amastigotes. **(A)** Serum lipid profile of 28 days LD-infected Balb/c mice (N=5). Equal number of sorted metacyclics promastigotes of two representative LD-S (LD-S^1^, LD-S^2^) and LD-R (LD-R^1^, LD-R^2^) strains were used to perform independent infection. **(B)** Number of attached LDs on KC surface 4Hrs p.i, data represents (N=3) for two LD-S and two LD-R strains. **C (i)** Representative Giemsa stain images of LD-S^1^ and LD-R^1^-infected KC at 4Hrs, 24Hrs and 48Hrs p.i Black arrow represents LD amastigotes. N denotes host nucleus. Scale bar 20μM. Giemsa Images are represented in grey scale to clearly represent LD nucleus (black arrow). **C (ii)** Intracellular amastigote count for 4Hrs, 24Hrs and 48Hrs p.i. Each dot represent count from 100 infected-KC (N=6). **D (i)** LD-infected-KC imaged to visualize the PV at 4Hrs and 24Hrs p.i. PV marked by ATP6V0D2 (magenta), KC and LD nucleus (blue), Scale bar 5μm. Yellow arrow represents LD amastigotes. In the merged image, white dotted line marks KC’s periphery, and host nucleus is represented with N. **D** (**ii)** PV counts measured from 30 infected KCs. **D (iii)** Confocal 3D reconstruction illustrating the spatial distribution of parasitophorous vacuoles (PVs) in Kupffer cells (KCs) infected for 24 hours. ATP6V0D2, a lysosomal vacuolar ATPase subunit, is visualized in magenta, while the nucleus is depicted in cyan. The final panel highlights PV structural grooves outlined in red solid lines, with intracellular *Leishmania donovani* (LD) amastigotes indicated by white arrows. Higher magnification of Figure 1D further emphasizes the increased abundance of PVs in LD-R infected cells, suggesting enhanced intracellular replication and adaptation mechanisms of drug-resistant strains. Scale bar 5µM. Both yellow and magenta solid line box represents the same area of the image. *** signifies p value < 0.0001, * signifies p value < 0.05, n.s non-significant.

### 2. Lipid deprivation is detrimental for LD-R while inherent hypercholesterolemia facilitates its proliferation

To determine if extracellular lipids are absolutely essential for proliferation of LD-R amastigotes, LD-S and LD-R-infections were performed in high, low or lipid-free medium. Reduction in extracellular lipids severely compromised proliferation of LD-R but not LD-S amastigotes in infected-KCs, confirming LD-R’s dependency on external lipid availability (**Figure 2A i, ii**). Alternatively, infection performed in KCs isolated from ApoE-knockout (KO)-mice(*Apoe^-/-^*) with inherent diet induced hyperlipidemia [32], resulted in significant increase in amastigote load as compared to wild type BL/6-infected-KCs, only for LD-R-infection, with LD-S infection resulting in comparable amastigote load (**Figure 2B i, ii**). Previous report has suggested that LD infection in hypercholesteremic Apoe^-/-^ mice triggers a heightened inflammatory response at approximately six weeks’ post-infection compared to wild type BL/6 mice, leading to more efficient parasite clearance. This is owing to unique membrane composition of Apoe^-/-^ which rectifies leishmania mediated defective antigen presentation at a later stage of LD infection [20]. Additionally, previous studies have also indicated that LD infection is well-established in mice within 6 to 11 days post-infection in murine models [33]. Thus to evaluate impact of initial lipid utilization on LD amastigote replication *in vivo*, BL/6 and diet-induced hypercholesterolemic Apoe^-/-^ mice were infected with GFP expressing LD-S or LD-R promastigotes and sacrificed 11 days p.i. Interestingly, however, we observed *Apoe^-/-^* mice infected with GFP expressing LD-S or LD-R resulted in increased splenic parasite burden as compared to wild type BL/6 mice at an earlier time point,11 days p.i (**Figure 2C i, ii**), This observation possibly indicates a higher initial infection propensity in *Apoe^-/-^* mice, which might get resolved later on due to increasing T-cell activity. Our observation also showed that splenic parasite load for LD-R is significantly higher in *Apoe^-/-^* than BL/6 (**Figure 2C ii, iii)** again confirming that higher availability of extracellular lipids might help in increased proliferation of LD-R amastigotes.

**Figure 2.**
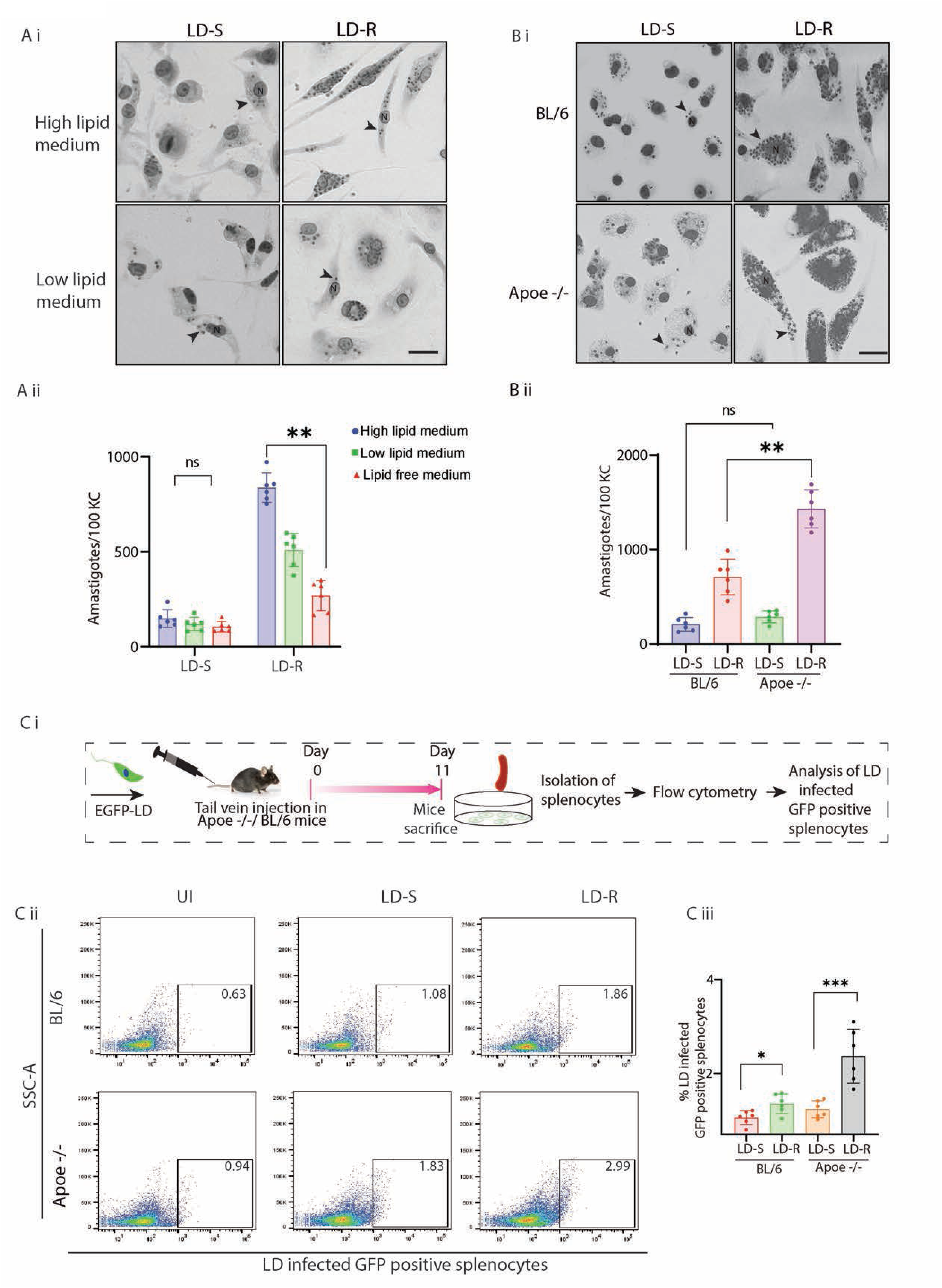
LD-R-infection results in a higher organ parasite load in hypercholesterolemic *Apoe^-/-^*mice. A. **(i)** Representative Giemsa images of LD-infected-KCs cultured in high and low lipid media. Black arrow represents LD amastigotes; N represents host nucleus. Scale bar 20μM. **A (ii)** Graph representing number of intracellular LD amastigotes in KCs with LD infection performed either in high, low or lipid free media (N=6). **B (i)** Giemsa images of LD-infected-KCs isolated from wild type BL/6 and *Apoe^-/-^* mice. Scale bar 20μm, N representing host nucleus. Giemsa Images are represented in grey scale to clearly represent LD nucleus (black arrow). **B (ii)** Amastigote count from 100 LD-infected-KCs (N=6) as in B(i). Data are presented as mean ± SEM. **C (i)** Schematic representation of *in-vivo* LD-infection in BL/6 and *Apoe^-/-^* mice performed with EGFP-LD-S or EGFP-LD-R **C (ii)** Flow cytometry representing GFP-positive splenocytes isolated from 11 days EGFP-LD infected BL/6 and *Apoe^-/-^* mice. The black box indicates % of GFP-positive splenocytes (LD-infected). **C** (**iii)** Graph representing data from six infected mice as in C (ii) are presented. *** signifies p value < 0.0001, ** signifies p value <0.001, * signifies p value < 0.05, n.s non-significant.

### 3. Quenching of host membrane cholesterol remains consistent during initial infection with LD-S and LD-R

Host lipids scavenging is essential for any intracellular pathogens exhibiting lipid auxotrophy [22]. Thus, to support their aggressive division, LD-R amastigotes should require significantly higher amount of host lipids, which in turn will allow excess supply of cholesterol for increased PV and daughter membrane biogenesis as compared to LD-S amastigotes. *Leishmania* amastigotes are lipid auxotrophs [11, 14], and it has been previously reported that *L. Mexicana* can acquire host membrane cholesterol during its initial entry into host MFs [34]. Thus, host membrane Cholesterol was investigated as a possible source and potential mechanism by which LD-R acquires host-derived lipids distinctively from their LD-S counterparts. However, confocal image analysis of NBD-Cholesterol labeled KCs infected with RFP expressing LD-S or LD-R revealed that both LD-S and LD-R can extract NBD-Cholesterol from the KC membrane during their initial infection (4Hrs p.i) with comparable efficiency (**Figure S2A i, ii**). This observation was further confirmed by quantification of NBD-Cholesterol between freshly invaded LD-S and LD-R parasites isolated from infected-KC by flow cytometry (**Figure S2B i, ii**).

### 4. LD-R-infection mobilizes extracellular LDL within infected-KCs to support its intracellular proliferation

Since the quenching of host membrane cholesterol during initial entry of LD promastigotes does not significantly differ in LD-S and LD-R mediated infections (**Figure S2**), we explored other potential host lipid sources for supporting LD-R’s increased intracellular division and membrane synthesis. LDL serves as the primary source of cellular lipids which helps to maintain lipid homeostasis, and gets readily metabolized to produce bioavailable cholesterol [35]. Our previous observation showing a significant drop in LDL levels between LD-S and LD-R-infected mice (**Figure 1**), led us to investigate LDL as a possible source which can differentially contribute in providing excess lipids, sterols, and fatty acids to rapidly proliferating intracellular LD-R amastigotes. Live cell imaging comparing uptake of fluorescent red-LDL in EGFP-LD-S or EGFP-LD-R-infected KCs at 24Hrs p.i revealed a significantly higher red-LDL accumulation within EGFP-positive LD-R-infected KCs as compared to LD-S-infection (**Figure 3Ai**). Fluorescence plate reader-based quantification further supported this observation showing significantly higher LDL accumulation in LD-R-infected-KCs over time, which was not observed by performing infection with killed parasites (**Figure 3A ii**), indicating that, LDL-influx is mediated by replicating LD amastigotes. Confocal microscopic images at 24Hrs p.i. of LD-S and LD-R-infected-KCs confirmed significantly higher LDL co-localizing with LD-R amastigotes compared to LD-S amastigotes (**Figure 3B i, ii**).

**Figure 3.**
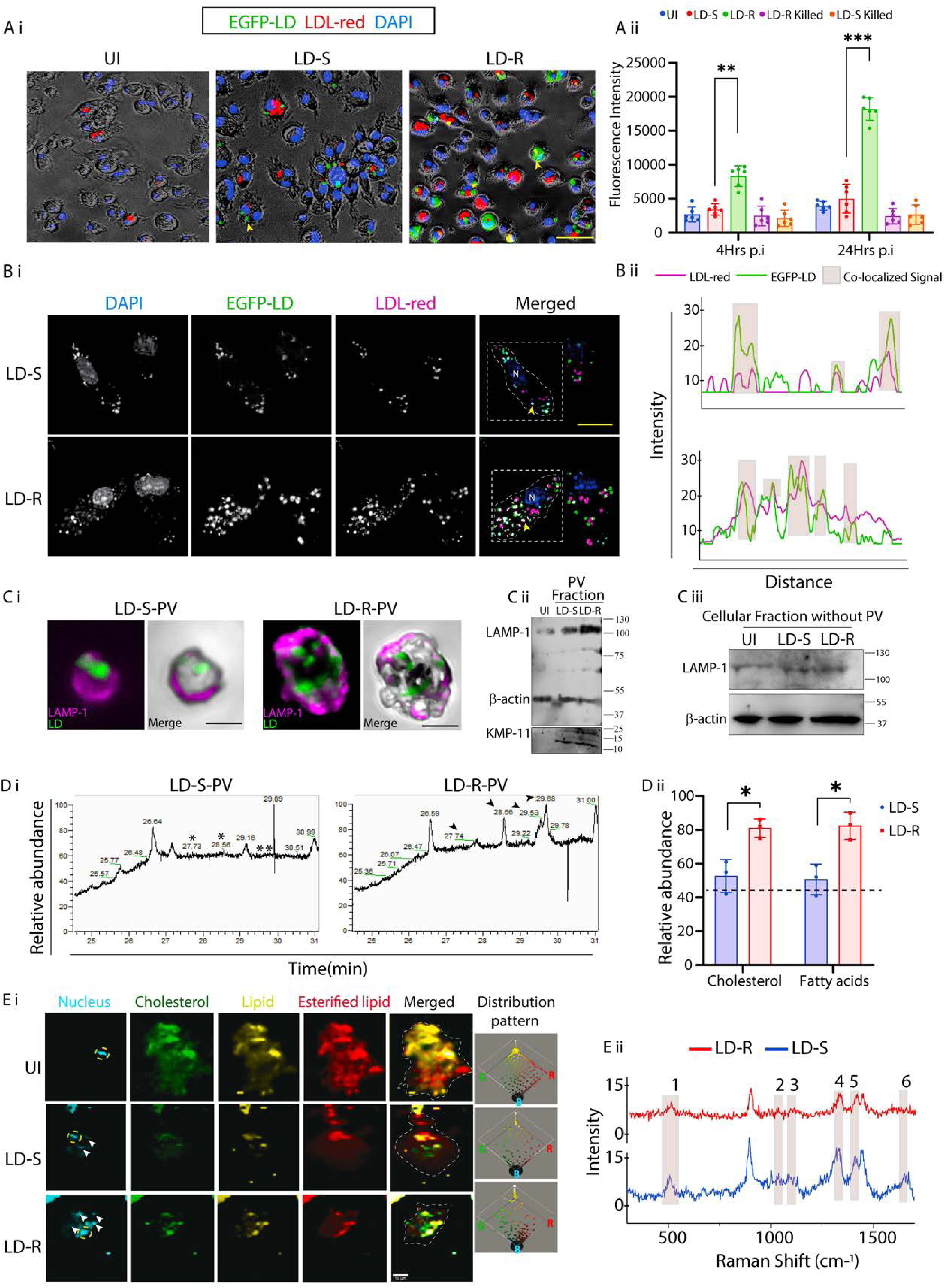
Endocytosed-LDL is the primary lipid source for intracellular LD-R proliferation. **A (i)** Live cell images of LDL-endocytosis in 24Hrs post infected KCs with LDL (red), LD (green), and the nucleus (blue). Images were acquired directly from 96 well glass bottom plate and bright field was merged with reduced brightness and increased contrast identically for all experimental conditions. Scale bar 40μm, Yellow arrow represents intracellular LD. **A (ii)** Graph representing Fluorescence plate reader-based quantification of A (i). **B (i)** Co-localization of LDL with the LD-amastigotes in infected KCs 24Hrs p.i. Merged image showing LDL (magenta), LD (green), nucleus (blue), White dotted line in the merged image demarcates infected-KC for co-localization analysis. Scale bar 10μm, yellow arrow represents LD nucleus and N represents host nucleus **B (ii)** Intensity plot representing co-localized signals from B (i), marked with grey transparent box. **C (i)** Confocal image showing the isolated LD-S and LD-R-PV, stained with LAMP1 (magenta) and LD (Green). Scale bar 2µm **C (ii)** Western blot analysis validated the purification of PV fractions, indicated by LAMP-1 positivity, while the presence of LD parasites within the PV was confirmed by KMP-11 detection**. C (iii)** Western blot of cellular fraction without PV showing minimal presence of LAMP1. **D (i)**GC-MS lipid profile of isolated PVs from LD-infected-KCs. Retention time 27.73 corresponding to the cholesterol and 28.56 corresponding to fatty acids are represented with * for LD-S-PV and with black arrowheads for LD-R-PV. **D (ii)** Relative abundance of Cholesterol and Fatty acids of D (i). **E (i)** Representative Confocal Raman spectroscopy image of uninfected or LD-infected-KCs 24Hrs p.i, illustrating distinct lipid-related signal distribution patterns marked with pseudo-colours. Yellow dotted circle in the left most panel demarcates the host cell nucleus while white arrow marking intracellular LD nucleus. In merged panel white dotted line demarcates cells periphery. The right most panel shows dot plot representation of lipid distribution, where each colour corresponds to different lipid-related peaks with respect to blue (B) indicating the nucleus. **E (ii)** Comparative Raman spectra from LD-S and LD-R-infected-KCs. Lipid-related peaks are demarcated with shaded box, representing 1. 540-560 cm-1 (Cholesterol), 2. 1080-1090 cm-1 (phospholipids), 3. 1270-1280 cm-1 (triglycerides), 4. 1300-1340 cm-1 (Amide-III bond), 5. 1440-1453 cm-1 (Fatty acids and Triglycerides), and 6. 1650-1660 cm-1 (Amide-I bond). *** signifies p value < 0.0001, ** signifies p value <0.001, * signifies p value < 0.05, n.s non-significant.

### 5. LD-R-infection results in a distinct metabolic shift in lipid peaks of infected host

Next, we wanted to confirm if higher LDL-uptake could result in accumulation of excess lipids, particularly cholesterol, in LD-R-PV that would provide LD-R amastigotes with higher replicative capabilities and associated enhanced membrane synthesis potential. Parasitophorous vacuole fractions were isolated from LD-S and LD-R-infected KCs at 24Hrs p.i. using a previously established protocol [36]. Following isolation, PV purity was confirmed through LAMP-1 staining which showed a significant enrichment around isolated PV in Confocal microscopy (**Figure 3C i**). Purity of isolated PV fractions was further confirmed by Western blot which showed an enhanced enrichment of LAMP-1 for LD-R-PV fraction as compared to LD-S-PV fraction, while PV excluded cellular fraction showed residual LAMP-1 expression confirming the purity of the isolated PV fractions **(Figure 3C ii, iii).** Following isolation, protein concentration was measured for isolated PV fractions using the Bradford assay, and PV fractions from both LD-S- and LD-R-infected KCs were normalized accordingly. Quantitative analysis of total cholesterol using Amplex red kit (**Figure S3A)** and thin layer chromatography (TLC) (**Figure S3B**) revealed significantly higher cholesterol content in LD-R-PV as compared to LD-S-PV. Furthermore, GC-MS of LD-S and LD-R-PV under identical experimental condition identified prominent peaks for cholesterol and fatty acids in LD-R-PV which were missing for LD-S-PV (**Figure 3D i, ii**). Apart from this biochemical characterization, we also performed biophysical analysis of lipid related peaks using non-invasive Confocal Raman spectroscopy to capture a snapshot of lipid distribution in LD-S and LD-R-infected-KCs under native conditions (**Figure 3E i, ii**). This analysis highlighted a distinct clustering of lipid peaks representing cholesterol, triglycerides, and fatty acids [37] around LD-R amastigotes, contrasting with their minimal clustering around LD-S amastigotes which resemble uninfected-KCs more closely (**Figure 3E i**). Raman Shift analysis within the biologically active region (300cm^-1^-1800cm^-1^) [38], further confirmed a significant difference in relative abundance for lipid related peaks’ at wavelengths: 540-560 cm^-1^ (cholesterol), 1080-1090 cm^-1^ (phospholipids), 1270-1280 cm^-1^ (triglycerides), 1300-1340 cm^-1^ (amide-III bond), 1440-1453 cm^-1^ (fatty acids and triglycerides) [37, 39], along with olefinic stretch of C=C double bonds at 1,655 cm^-1^ between LD-S and LD-R-infected-KC, at 24Hrs p.i. **(Figure 3E ii and S3C i, ii**). We also noticed a distinct shift by plotting area under the curve from (Right to Left) in response to LD-R-infection (**Figure S3 D**), indicative of distinctly different lipidomes [40].

### 6. LDL-uptake in LD-infected-MΦs occurs through fluid phase endocytosis

As LD-R amastigotes heavy reliance on increased host lipid acquisition for rapid proliferation became evident, we wanted to investigate molecular basis of this acquisition. Considering that MFs primarily uptake LDL via the LDL receptor (LDLr) [41], we checked its status in response to LD-S and LD-R-infection. However, immunofluorescence analysis showed a decrease in LDLr expression in infected-KCs harboring LD-amastigotes as compared to adjacent uninfected KCs, regardless of infection type (**Figure 4A**), along with no significant changes in total LDLr expression between uninfected, LD-S, and LD-R-infected-KCs, as observed from Western blot (**Figure S4A**). This indicated that LDL-uptake by LD-infected-KCs might be receptor-independent. Our attempts to generate siRNA mediated Knock down (KD) in KCs were unsuccessful, hence to convincingly determine the role of LDLr in LD-infected-MFs we generated LDLr-KD-PEC (**Figure S4B i, ii**) and performed infection with LD-S or LD-R. Infecting LDLr-KD-PEC with LD-S or LD-R resulted in no significant changes in intracellular amastigote numbers for both infection types as compared to scrambled RNA control (**Figure SC**) confirming that although LD-R relies on increased LDL-influx for heightened proliferation, this influx is LDLr-independent. Apart from LDLr, MFs can also acquire LDL through receptor-independent fluid phase endocytosis [42]. To confirm this possibility, we checked the status of Cofilin, an actin modulating protein, which serves as a marker for fluid phase endocytosis. Western blot analysis of infected-KCs revealed an increase in Cofilin levels with a concurrent decrease in its phosphorylated form as LD-R-infection progressed, indicating active fluid phase endocytosis (**Figure 4Bi, ii**). In contrast, LD-S-infection did not induce such changes (**Figure 4B i, ii**). Interestingly, this decrease in Cofilin phosphorylation was also observed in LD-R-infected-PEC (**Figure S4D i, ii**), signifying that enhanced fluid phase endocytosis may be a conserved mechanism for facilitating LDL-influx in LD-R-infected-MFs. This enhanced fluid phase endocytosis in response to LD-R-infection was further confirmed by structural illumination imaging (SIM), which showed increased Cofilin turnover with dispersed distribution in LD-R-infected-KCs, leading to enhanced F-Actin rearrangement and protrusion formation [43], which was absent for uninfected or LD-S-infected-KCs (**Figure 4Ci, ii and Figure 4Di, ii**). Finally, we infected the KCs with GFP expressing LD-R for 4Hrs, washed and allowed the infection to proceed in presence of fluorescent red-LDL and Latrunculin-A (5µM), a compound which specifically inhibits fluid phase endocytosis by inducing actin depolymerization [42]. Real-time fluorescence tracking demonstrated that Latrunculin-A treatment not only prevented the uptake of fluorescent red-LDL but also severely impacted intracellular proliferation of LD-R amastigotes (**Movie 3A and 3B and Figure 4E**). In contrast, treatment with Cytochalasin-D, which alters cellular F-actin organization but does not affect fluid phase endocytosis [42], had no effect on the intracellular proliferation of LD-R irrespective of Cytochalasin-D concentrations (2.5µg/ml and 5µg/ml respectively) (**Figure 4F and Figure S4E**).

**Figure 4.**
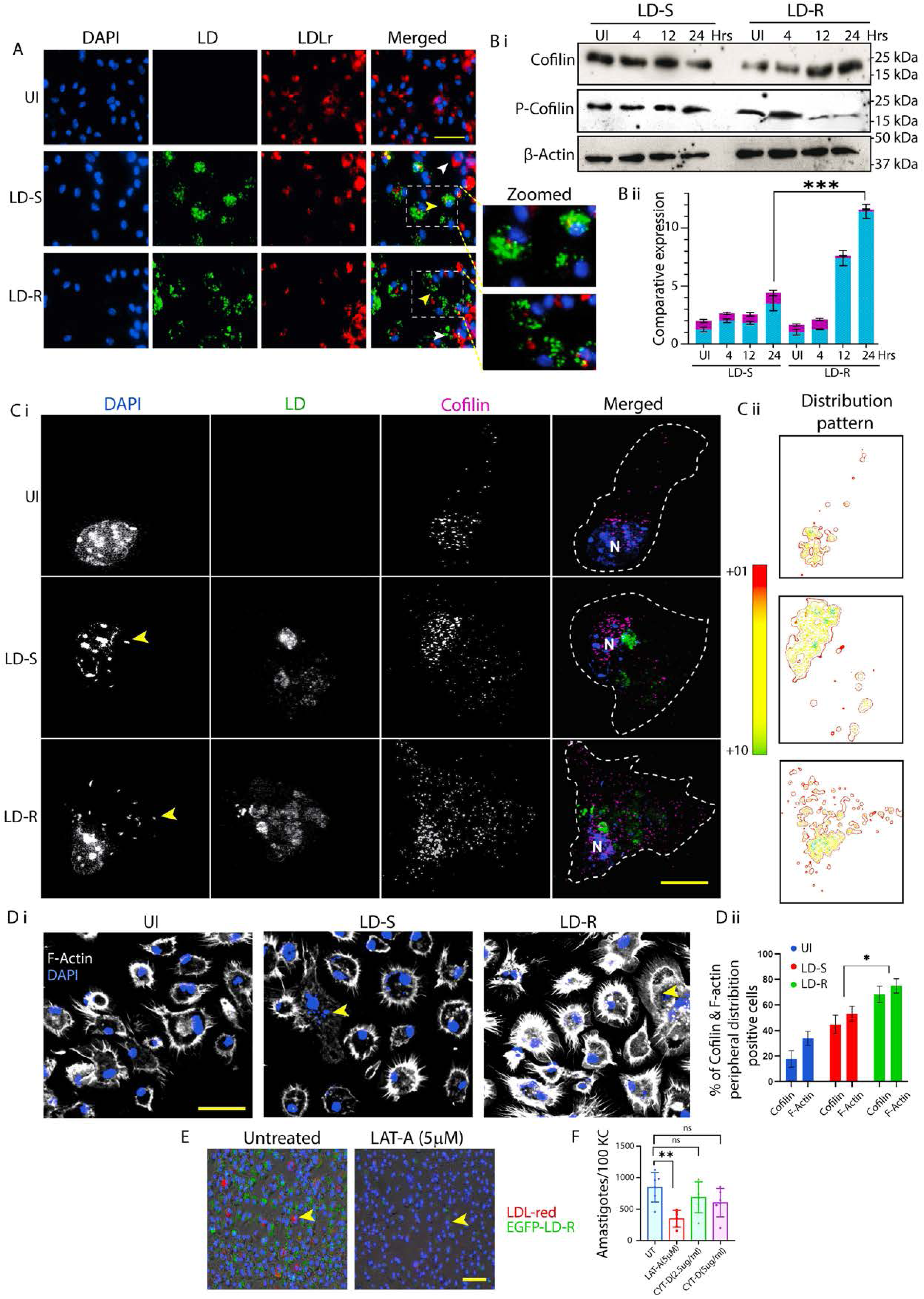
Increased LDL uptake in LD-R-Infected-MFs through receptor independent fluid phase endocytosis. **(A)** Immunofluorescence images of LDLr expression in LD-infected-KCs. Scale bar 40μm. Enlarged view of A showing infected-KC with low LDLr expression (yellow arrow) compared to neighboring uninfected KC (white arrow) in the merged image**. B (i)** Cofilin and phosphorylated-Cofilin expression by western blot in LD-infected**-**KCs. **B (ii)** Graphical representation of comparative expression of Cofilin and phosphorylated-Cofilin with respect to β-actin for B (i). **C (i)** SIM images showing the Cofilin distribution in LD-infected-KCs. Yellow arrow represents intracellular LD, white dotted line in the merged image demarcates cell boundary. Scale bar 5μm. **C (ii)** Colour spectrum representing Cofilin distribution pattern of C (i) with green representing maximum and red representing minimum distribution. **D (i)** Confocal images representing Filamentous actin (F-actin) protrusions in LD-infected-KCs. Scale bar 20μm, with yellow arrow representing LD amastigotes. Individual experimental condition is represented in pseudo-colour, with uniform high contrast to clearly represent the F-actin protrusions. **D (ii)** Graph representing Cofilin and F-actin distribution in KCs infected with LD or left uninfected. (**E)** Still images from live cell video microscopy representing co-localization of LD-R with LDL in presence or absence of Latrunculin-A (LAT-A). Scale bar 60μm. Yellow arrow represents LD-infected-KCs with LDL(red). **(F)** Graphical representation (N=6) of amastigote count in 100 LD-R-infected-KCs treated with either LAT-A, or Cytochalasin (CYT-D) or left untreated. Data are presented as mean ± SEM. ** signifies p value <0.001, * signifies p value < 0.05, n.s non-significant.

### 7. Endocytosed-LDL traffics and fuses with and LD-R-PV with increased proficiency

Typically, LDL-vesicles fuses with lysosomes upon entry into MFs, leading to subsequent transfer of LDL-derived cholesterol into the endoplasmic reticulum and cytoplasm [44]. As we have already confirmed an increased influx with higher presence of endocytosed-LDL vesicles around LD-R amastigotes along with significantly higher accumulation of cholesterol and fatty acids in LD-R-PV as compared to LD-S-PV (**Figure 2**), we hypothesized that the mode of trafficking of endocytosed-LDL might differ between LD-S and LD-R-infection. LD-PV are modified phagolysosomes [36, 45], and we observed an increased convergence of LDL-vesicles with LAMP-1 positive LD-R-PV even at 4Hrs p.i, while such convergence was not noticed in LD-S-infected-KC (**Figure 5A i, ii**). Structured Illumination Microscopy confirmed a complete convergence of LDL-vesicles with LD-R-PV at 24Hrs p.i, while LD-S-PV showed only partial convergence with LDL-vesicles even at this late infection point (**Figure 5B i, ii)**. Interestingly, while, Western blot analysis revealed minimal expression of LAMP-1 in response to LD-S-infection at 4Hrs p.i, which increased considerably as the infection progressed (**Figure 5C i)**, however, a considerable amount of LAMP-1 expression was observed for LD-R-infected-KCs from 4Hrs p.i, which also increased as infection progresses (**Figure 5C ii)**, indicating that a higher fusion of LDL-vesicles in LD-R-PV may be accompanied by increased lysosomal biogenesis required for degrading LDL into cholesterol. Filipin staining of EGFP-LD-S and EGFP-LD-R-infected KCs at 24Hrs p.i. revealed enhanced cholesterol accumulation surrounding LD-R amastigotes, as indicated by a significantly higher co-localization coefficient compared to LD-S amastigotes (**Figure 5D i, ii, iii**). Importantly, this cholesterol enrichment was absent around dextran beads harboring LD-R amastigotes, suggesting that cholesterol accumulation is a process specifically associated with LD-R infection. Infected-KCs harboring both LD-R amastigotes and Fluorescent Latex Beads, showed a concentrated staining of cholesterol around LD-R amastigotes, with no positive Cholesterol staining around internalized latex beads similar to LD-S amastigotes (**Figure 5E)**.

**Figure 5.**
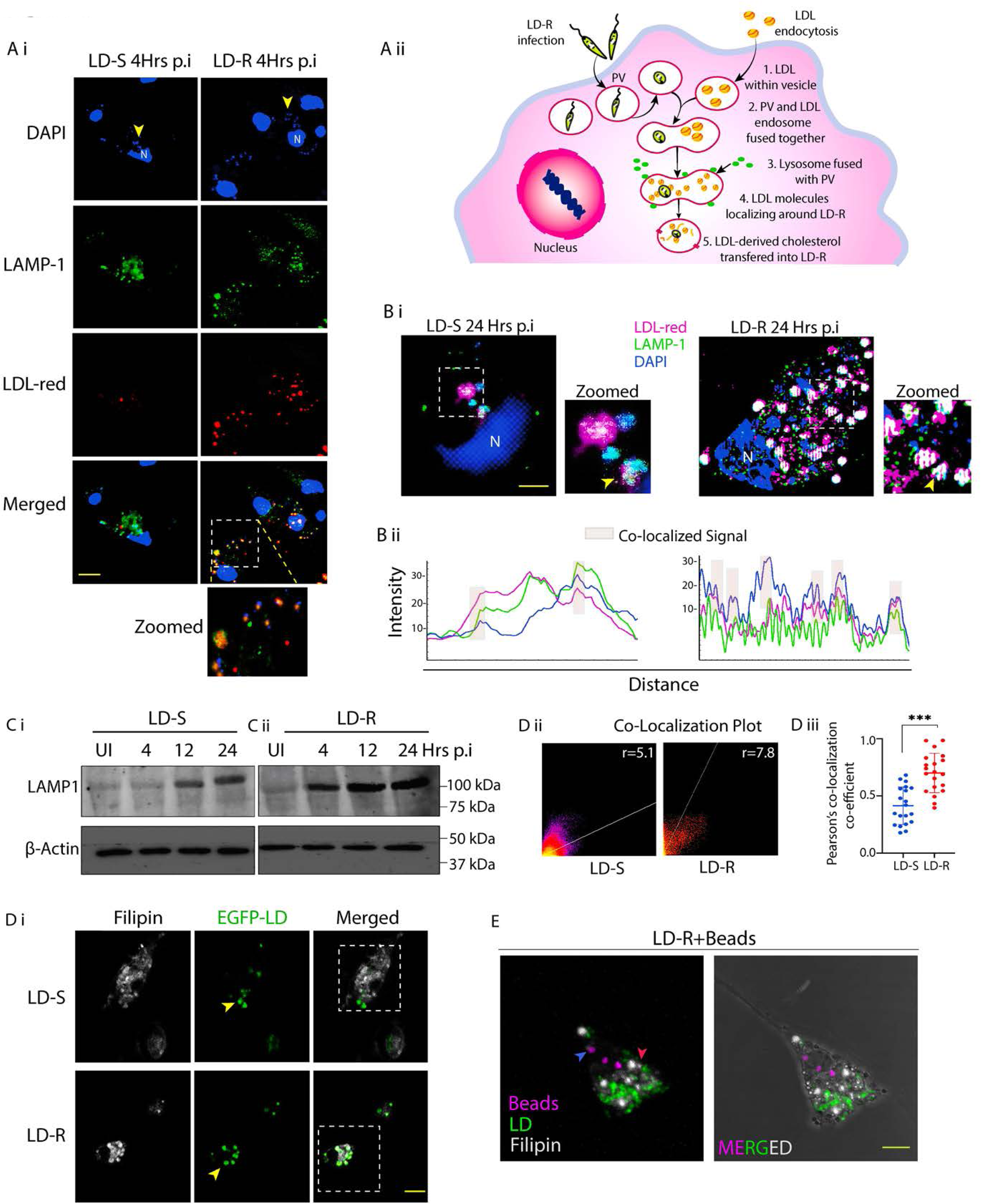
Endocytosed-LDL fuses with LD-R-PV to provide Cholesterol to LD-R amastigotes. **(A)** Confocal images of representing LDL-vesicles, with LAMP1 positive lysosomal vesicles in LD-infected-KCs 4Hrs p.i. Yellow arrow represents LD nucleus, white dotted box marks the region further cropped and zoomed to show fusion of lysosomal vesicles with LDL and LD-R. Scale bar 10μm. **A (ii)** Schematic representation showing trafficking of LDL-vesicle towards the LD-R-PV. **B (i)** SIM images representing convergence of LDL-vesicles with LD amastigotes and LAMP1 in 24Hrs with LD-infected-KCs. White dotted box marks the region further cropped and Zoomed to clearly show the convergence (of LD amastigotes with LDL and LAMP-1 (white vesicles shown by yellow arrow), Scale bar 5μm. **B (ii)** Intensity plot representing of co-localization signals for B (i). **C** Western blot analysis showing LAMP1 expression in LD-infected-KCs from early to late hours. **C (i), C (ii)** representing LAMP1 expression LD-S and LD-R infected-KC **D (i)** Filipin staining of EGFP-LD-infected-KCs. Filipin (white) and LD (green), yellow arrow represents intracellular LD. Scale bar 10μm. White dotted line marked cells used to generate co-localization plot. **D (ii)** Co-localization signals of LD (EGFP and Filipin represented by the dot plot. Pearson co-localization coefficient value (r) represented for LD-S and LD-R-infected-KC. **D (iii)** Graph representing Pearson’s coefficient for EGFP-LD-infected-KCs (N=19). **E** Filipin staining of EGFP-LD-R-infected KCs incubated with dextran beads (Magenta) revealed minimal colocalization between the beads and Filipin, while the LD-R amastigotes within same macrophage shows a significant co-localization with Filipin. Blue arrow demarking beads with limited filipin colocalization, red arrow demarking LD-R with filipin colocalization. Scale bar 5µM. Statistical significance in the observed differences was determined through ANOVA utilizing GraphPad Prism software (version 9.0). *** signifies p value < 0.0001.

### 8. LD-R-infection suppress release of metabolized cholesterol from PV

Once LDL is broken down to cholesterol by lysosomal acid hydrolases, metabolized cholesterol is released back into cytosol from LDL-vesicles by two transmembrane proteins namely NPC-1 and NPC-2 [44]. We suspected that LD-R must prevent or delay this process as this will limit cholesterol accessibility to replicating LD-R amastigotes. Analysis of available *RNAseq* (INRP000146) data for uninfected, LD-S and LD-R-infected-PEC (**Figure 6A)** revealed a suppression of *npc-1* expression along with its transcription factor *srebp2* [46], in response to LD-R-infection. Western Blot analysis of LD-S and LD-R-infected-KCs confirmed this low expression of SREBP2 and NPC-1 in LD-R-infected-KC 24Hrs p.i (**Figure 6B i, ii**). More interestingly, Confocal imaging not only confirmed this low expression of SREBP2, but even in those cases of LD-R-infected-KCs where we did observe some residual expression of SREBP2, it was entirely restricted to cytoplasm, unlike LD-S or uninfected-KCs where SREBP2 did get translocated in the nucleus (**Figure 6C**). Furthermore, as SREBP2, acts as the master regulator of cholesterol biosynthesis, its downregulation and inhibition of nuclear translocation further impacted expression of the downstream cholesterol biosynthetic enzyme HMGCR (**Figure 6D**) specifically in LD-R-infected KCs. This again supports the *RNAseq* expression data (**Figure 6A**) and further confirms that LD-R-infected KCs has a shutdown of *de novo* cholesterol synthesis. For NPC-1, immunofluorescence imaging comparing adjacent uninfected and infected-KCs harboring LD-S or LD-R, clearly confirmed unlike LD-S, LD-R specifically suppresses NPC-1 to prevent release of LDL-derived cholesterol from fused vesicles making it accessible to amastigotes (**Figure 6E**). We confirmed this observation by performing a Total Internal Reflection Microscopy (TIRF) which clearly showed a low plasma membrane cholesterol for LD-R-infected-KCs as compared to LD-S-infected-KCs (**Figure 6Fi**) and subsequently revalidated this by quantifying the cholesterol content in isolated membrane fraction for individual experimental set as mentioned above (**Figure 6F ii**). Finally, siRNA mediated knock down of NPC-1 in PEC (due to failure of successful KD generation in KCs), resulted in significant increase in amastigote load in response to LD-R-infection, while no such change in amastigote load was observed for LD-S-infected NPC-1-KD-PECs (**Figure 6G i, ii, iii, iv**).

**Figure 6.**
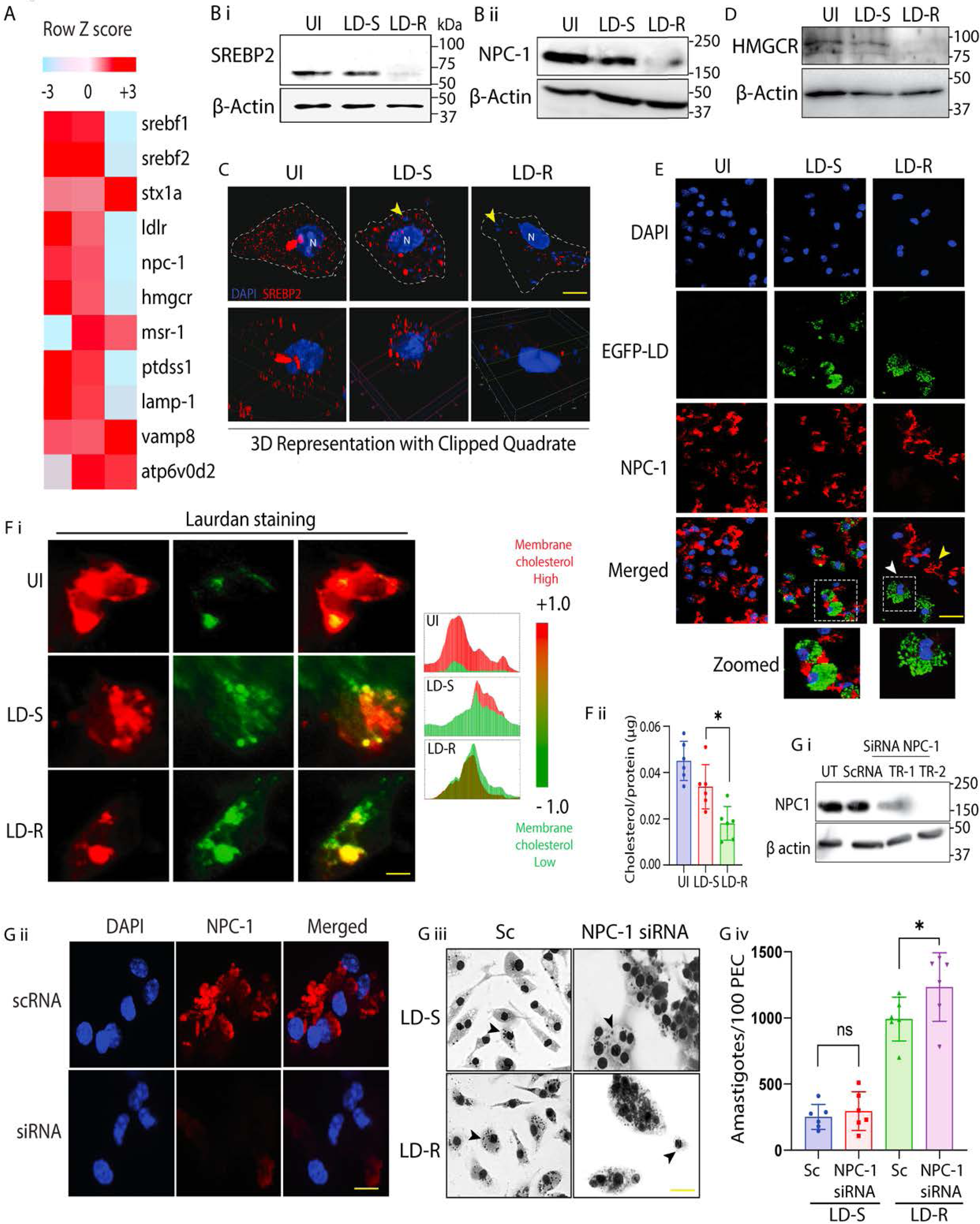
LD-R infection suppress NPC-1 to alter Cholesterol mobilization in infected MFs. **(A)** 24Hrs LD-S and LD-R-infected-PECs were subjected to *RNAseq* analysis keeping uninfected-PECs as control. Heat map representing average expression of differentially expressed genes related to lipid metabolism between LD-S and LD-R-infected-PEC. Scale represents median-centered counts (TPM). Row Z score data represented here. Expression of RNAs (right margins) is presented as centered and ‘‘scaled’’ [mean normalized log_2_(TPM + 1)]. **B (i)** Expression of SREBP2 and **B (ii)** NPC-1 in LD-infected and uninfected KCs by Western blot with β-actin as control. **C** Intracellular localization of SREBP2 in response to LD-infection in KC 24Hrs p.i. White dotted line represents cell periphery with yellow arrow representing intracellular LD and N representing host nucleus. 3D image represented below with one quadrant clipped to confirm nuclear translocation of SREBP2. Scale bar 5μm. **(D)** Expression of HMGCR LD-infected and uninfected KCs by Western blot with β-actin as control. **E** Immunofluorescence showing NPC-1 expression in LD-infected-KCs. White arrow representing NPC-1 expression in LD-R-infected-KCs and yellow arrow representing NPC-1 in adjacent uninfected-KCs in the merged panel. White dotted box marks regions further cropped and shown in the bottom panel with enlarged view (Zoomed). Scale bar 20μm. **F (i)** TIRF microscopic images of Laurdan stained LD-infected and uninfected KCs with two different spectra (488nm, 594nm), changes from red to green represents gel to fluid phase transition of the host plasma membrane. Colour distribution pattern corresponding to the TIRF images representing each experimental condition is presented in the right most panel. Scale bar 5μm. **F (ii)** Measurement of membrane cholesterol by Amplex red assay kit. Graphical representation showing total membrane cholesterol divided by total membrane protein. **G (i)** Expression of NPC-1 determined by western blot in PECs transfected with NPC-1 siRNA or scrambled (sc) siRNA. **G (ii)** Expression of NPC-1 determined by immunofluorescence in PECs transfected with NPC-1 siRNA or scrambled (sc) siRNA. Scale bar 5μm. **G (iii)** Giemsa-stained images of LD-infected NPC-1 knockdown-PECs with scrambled siRNA control (Sc). Scale bar 20μm. Images are represented in grey scale with increased contrast to represent LD nucleus (black arrow). **G (iv)** Graphical representation of number of intracellular LD-amastigotes as in F (ii). Results show the counting of 100 infected-PEC from 6 independent replicate. * signifies p value < 0.05, n.s non-significant.

### 9. LD-R-Infection specifically excludes oxidized-LDL in infected murine liver to limit inflammation

The increased accumulation of LDL within MFs typically triggers an inflammatory response [47]. This possessed an inherent conceptual dilemma to our observation as increased inflammatory response should be detrimental for LD-R amastigotes [48, 49]. To check this, we measured inflammatory IFN-γ by co-culturing splenic T-cells isolated from LD-S or LD-R-infected mice, with LD-S and LD-R-infected-KCs, respectively (**Figure 7A**). Surprisingly, our ELISA results showed a low IFN-γ levels in the culture supernatant of LD-R-infected-KCs even after 24Hrs of co-culture, while for LD-S-infection a significant increase was noted at this point (**Figure 7Bi**). Similarly, stimulation of LD-infected splenocytes obtained from LD-S and LD-R-infected *Apoe^-/-^* mice with specific soluble leishmamia antigens (SLA) of LD-S or LD-R also confirmed this restricted IFN-γ response to LD-R-infection (**Figure 7B ii)**, thus suggesting that although LD-R-infection results in increased accumulation of lipids in host MFs, it elicits a suppressed inflammatory response as compared to LD-S. Notably, accumulation of oxidized-LDL (ox-LDL), and not its neutral form has been reported to cause inflammatory response [47, 50]. LD-S or LD-R-infected-KCs incubated with ox-LDL showed a significant purple positive staining of ox-LDL only for LD-S-infected-KCs (**Figure 7C i**), with LD-R-infected-KCs seemingly inhibiting uptake of ox-LDL. Infecting KCs with higher number of LD-S or lower number of LD-R did not alter their ox-LDL uptake capabilities, clearly suggesting LD-R-infection inherently suppress ox-LDL uptake, irrespective of the number of intracellular amastigotes (**Figure 7C. ii**,). Although, receptor of ox-LDL, MSR-1 [26], have shown a slightly lower expression in LD-R-infected-PEC in the *RNAseq* analysis (**Figure 6A**), however owing to reported dominant expression of *msr-1* in the liver [50], we checked expression of MSR-1 in the LD-S and LD-R-infected-KCs. Confocal images and Western blot analysis confirmed a significant lower expression of MSR-1 in LD-R-infected-KCs as compared to LD-S-infected-KCs (**Figure 7D i, ii**), thus explaining reduced staining for ox-LDL in response to LD-R infection. Furthermore, increased expression of *atp6vod2* transcripts, implicated in LD-PV acidification and lipid utilization [36], was also noted to be slightly higher for LD-R-infected-PECs in the *RNAseq* (**Figure 6A**). However, western blot analysis showed a slightly increased expression of ATP6V0D2 in LD-R-infected-KCs as compared to LD-S-infected-KCs (**Figure 7E**), suggesting that it might also play a role in maintaining LD-PV acidity, facilitating lipid utilization and limiting ox-LDL uptake, as previously reported for *L. mexicana* infection [36]. Finally, immunostaining of liver sections from 28 days LD-S and LD-R-infected-mice **(Figure 7F i)** confirmed a significant lower expression of MSR-1 in LD-R-infected liver with higher accumulation of neutral lipid droplets as compared to LD-S-infected murine liver **(Figure 7F ii, iii),** thus confirming LD-R specifically accumulates neutral lipids by restricting MSR-1 expression.

**Figure 7.**
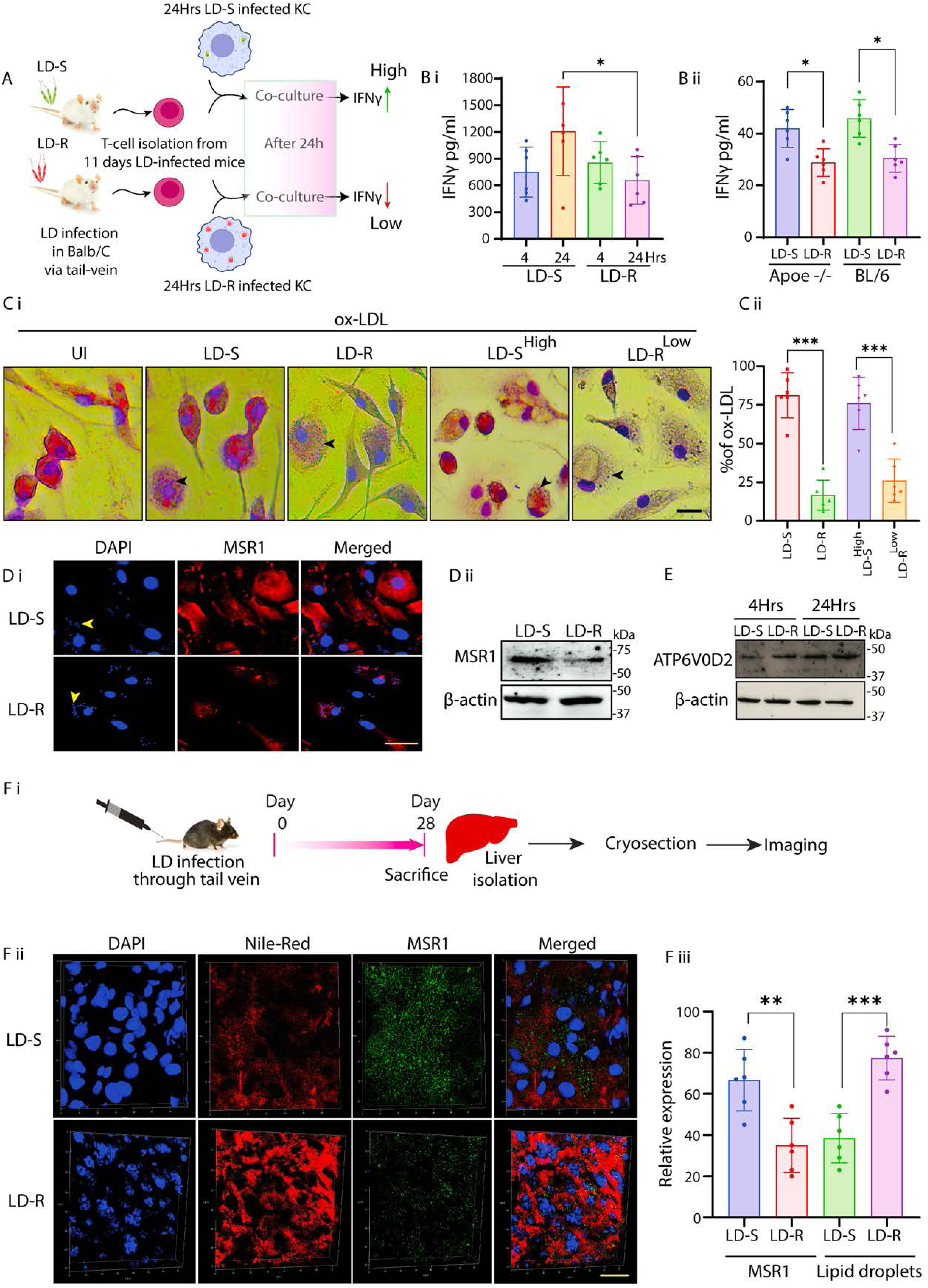
LD-R-infected-MFs selectively excludes ox-LDL to suppress inflammatory response in host. **(A)** Scheme representing *ex vivo* co-culture experiment performed by isolating T-cells from LD-infected mice with in vitro LD-infected-KCs **B (i)** IFN-**γ** level in the supernatant of LD-infected-KCs co-cultured with the T-cells isolated from spleen of Balb/c mice infected with LD-S or LD-R 28 days p.i. Data represented as mean ± SD from three independent experiments. **B (ii)** IFN-**γ** level in the supernatant of splenocytes cultures from LD-infected-BL/6 and *Apoe^-/-^*-mice stimulated with SLA (Soluble *Leishmania* antigen) specific for LD-S or LD-R. Data are presented as mean ± SD from six independent mice. **C (i)** ox-LDL incubated LD-infected-KCs was fixed and stained with Oil-red-O and Haematoxylin. LD-S^High^ represents MOI (1:20) and LD-R^Low^ represents MOI (1:5). Scale bar 10μm. Black arrow represents LD nucleus. **C (ii)** Graph representing % of ox-LDL positive MFs under different experimental conditions as represented in C (i). **D (i**) Confocal image showing MSR-1 in LD-infected-KCs. Yellow arrow represents LD-amastigotes while N represents host nucleus. Scale bar 20μm **D (ii)** Expression of MSR1 detected by Western blot of LD-infected-KC with β-actin as loading control. (**E)** Expression of ATP6V0D2 in LD-infected-KCs by Western blot with β-actin as loading control. **F (i)** Scheme representing the time line of *in vivo* experiment for IFA**. F (ii)** Expression of MSR1 and neutral lipid droplet accumulation assayed by Confocal microscopy from the cryo-sectioned liver of 28 days LD-infected-mice. Representative image showing Lipid droplet marked in red and MSR1 in green, nucleus in blue. Scale bar 40μm. **F (iii)** Graphical representation of % positive Lipid droplets accumulation and MSR-1 expression under different experimental condition as in E (i). Data collected from N=6 mice. *** signifies p value < 0.0001, ** signifies p value <0.001, * signifies p value < 0.05.

### 10. Lipid droplets accumulation decreases Amphotericin-B responsiveness in LD-R amastigotes

Increased LDL-uptake has been reported to cause high lipid droplets formation [51, 52], and lipid droplets formation has been reported to play critical role in intracellular proliferation of several pathogens, including LD-amastigotes [53–55]. As we have already observed significant accumulation of lipid droplets in LD-R-infected murine liver (**Figure 7**), we further investigated its role in response to LD-S and LD-R-infection. Lipid droplets staining of LD-S and LD-R-infected-KCs revealed significantly higher lipid droplets accumulation around LD-R amastigotes (**Figure S5A** and **Movie 4).** For better visualization, we expanded [56] EGFP-LD-R-infected-KCs and again performed lipid droplets staining on them which confirmed assimilation of neutral lipid into LD-R amastigotes (**Figure 8A**). This was further confirmed by performing TEM in LD-R-infected-KC which showed Lipid droplets accumulating around LD-R amastigotes **(Figure 8B)**. Lipid accumulation has been previously reported to cause Amphotericin-B unresponsiveness among pathogens by hindering drug penetration [23]. Interestingly, we noted that two LD-R-strains (BHU575 and BHU814) which we have included in this study also shows reduced responsiveness against Amp-B, a second-line-antileishmanial [57], and we validate this by determining their EC50 against Amp-B in LD-infected-KCs, which showed an EC50 of 0.39 μM and 0.31 μM for LD-R^1^ and LD-R^2^ amastigotes respectively, as compared to EC50 of 0.03 μM for LD-S^1^ and 0.032 μM LD-S^2^. We suspected that significant lipid droplets accumulation around LD-R amastigotes might be responsible for their reduced susceptibility towards Amp-B by limiting drug accessibility. To test this possibility, we performed independent infection with two LD-R strains in low lipid and high lipid medium and access their EC50 against Amp-B. Low lipid medium results in significant drop in lipid droplets accumulation within LD-R-infected-KCs (**Figure S5B**) and also resulted in enhanced susceptibility for both the LD-R strains against Amp-B with EC50 reduced to 0.39 μM to 0.16 μM for LD-R^1^ and 0.31 μM to 0.06 μM for LD-R^2^ as compared to high lipid medium (**Figure 8C i, ii)**. Interestingly, infection with lab-generated Amphotericin-B-unresponsive LD promastigotes (LD-S^Amp-B-R^) without primary unresponsiveness towards antimony did not result in any significant lipid droplets accumulation around LD-R amastigotes (**Figure S5A**) and neither showed any significant alteration in EC50 between low (0.09 μM) and high lipid medium (0.11 μM) (**Figure 8C iii**). This observation clearly suggests that this high LDL-influx along with higher lipid droplet accumulation responsible for increased unresponsiveness of LD-R isolates towards Amp-B is a specific attribute of LD isolates with primary unresponsiveness against antimony and may vary significantly from lab generated Amp-B unresponsive strains. Previously, Aspirin (Asp), a lipid droplet blocker has been reported to block lipid droplet formation in *T.cruzi* and LD-infected-MFs [53, 58]. We also observed a drastic drop in lipid droplets formation in LD-R-infected-KCs treated with Aspirin as compared untreated LD-R-infected-KCs (**Figure S5C).** Interestingly, while treatment with Aspirin alone did not significantly reduce the amastigote load in LD-R-infected-KCs, combining Aspirin with previously determined low dose of Amp-B (0.1 μM), completely restored Amp-B responsiveness against intracellular LD-R amastigotes (**Figure 8E, F**). This suggests that the enhanced uptake of LDL leading to lipid droplet accumulation might be a primarily responsible for increasing Amp-B unresponsiveness among clinical LD-R isolates. Importantly, this observation is consistent with findings from human VL patients from the endemic region of Bihar, India, where those who relapsed after Amp-B treatment had significantly lower serum LDL and cholesterol levels compared to Amp-B treatment responders (**Figure 8G and Table 2**).

**Figure 8.**
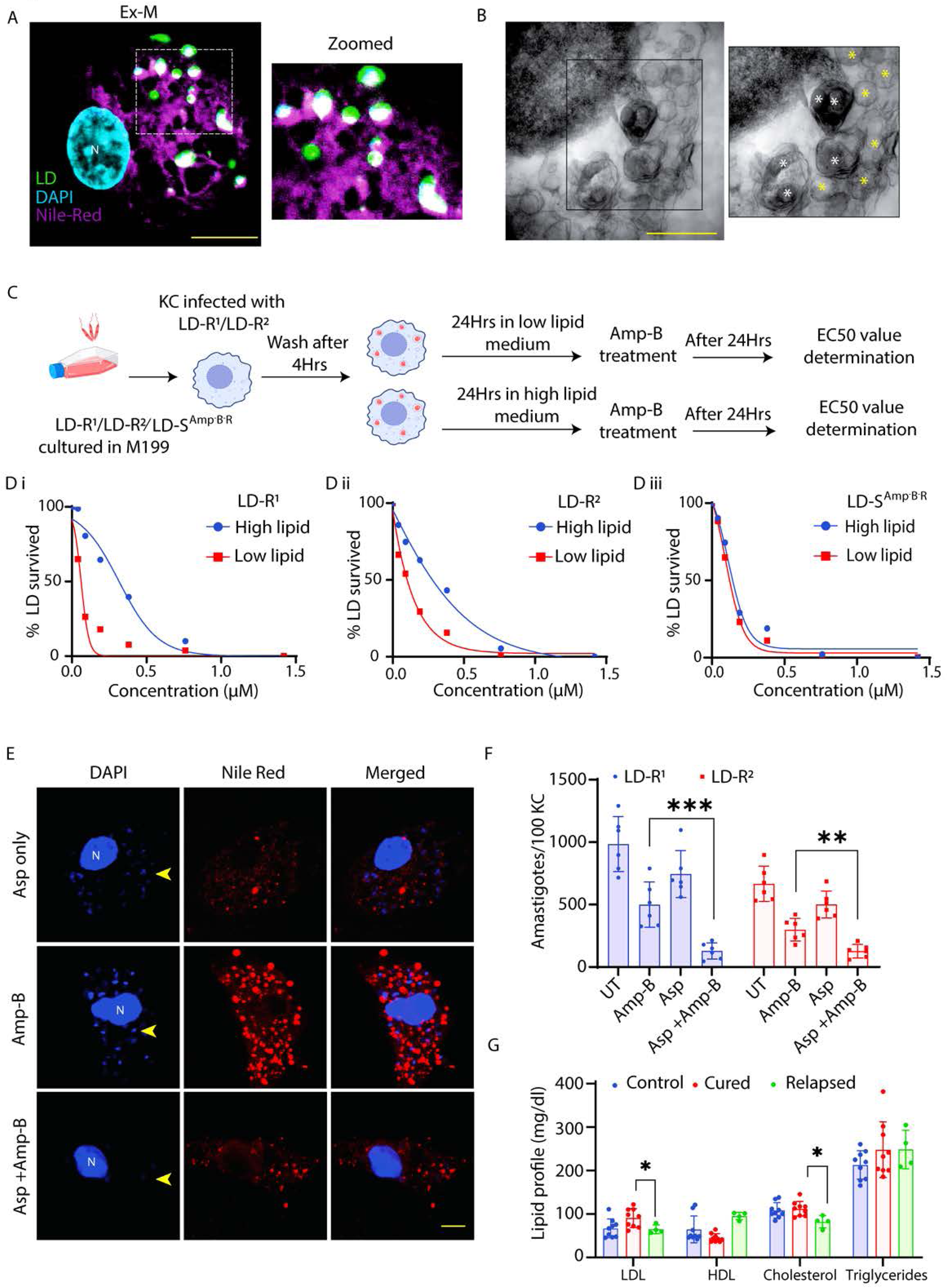
Lipid droplets accumulation around LD amastigotes is inversely related with susceptibility towards Amphotericin-B. **(A)** Ex-M image showing lipid droplet assimilation in LD-R-amastigotes. White dotted region further cropped and shown in enlarged panel (Zoomed). Nile red (magenta), LD (green), nucleus (cyan). White spots represent Lipid droplets assimilated in LD-R and Host nucleus is represented as N. Scale bar 5μm. **(B)** TEM images showing lipid droplets around the LD-R amastigotes. Black box further magnified and represented in right panel. White * representing LD-R amastigotes, yellow * showing lipid droplets. Scale bar 2μm. (**C)** Schematic representation representing experimental scheme used to determine EC50 against Amphotericin-B under different experimental conditions**. D (i) and (ii)** Determination of EC50 value against Amphotericin-B in low lipid and high lipid media by performing infection with of two independent LD-R strains. **D (iii)** Determination of EC50 value against Amphotericin-B in KC infected with LD-S^Amp-B-R^. Intracellular amastigotes were counted through Giemsa staining. **(E)** Confocal images showing lipid droplets accumulation around intracellular amastigotes with Aspirin (5 μM) or Amp-B (0.36 μM) or with a combination of Aspirin (5μM) and Amp-B (0.11μM) at 48Hrs p.i **(F)** Determination of amastigote load in KCs infected with two independent LD-R isolates in different experimental conditions either untreated or treated with Aspirin (5 μM) or Amp-B (0.36 μM) or with a combination of Aspirin (5μM) and Amp-B (0.11μM) at 48Hrs p.i (**G)** Comparative Lipid profile of VL patients (Cured and Relapsed) in response to Amp-B treatment along with healthy individuals from endemic region of Bihar, India. *** signifies p value < 0.0001, ** signifies p value <0.001, * signifies p value < 0.05.

## Discussion

Our findings revolve around understanding how antimony-resistant *Leishmania donovani* (LD-R) has evolved as a persistent strain by adopting to a strategy of better utilization of host derived lipids to sustain their own aggressive intracellular proliferation. Intracellular pathogens often alter the host’s immune-metabolic state for better survival [59, 60]. Our observations pointed out that high organ parasite load linked with LD-R-infection does not rely only on initial infectivity as commonly believed to date [7], rather is indicative of a selective higher intracellular proliferation in host. We observed LD-R amastigotes scrutinize and access specific type of host lipoproteins to support their aggressive proliferation as seen in clinical infection (**Figure 1, S1**) [15]. Previously increased amastigote load in VL patients has been inversely linked with their serum cholesterol level [29], and we also observed that mice infected with LD-R isolates had significantly lower HDL, LDL, and cholesterol compared to LD-S-infected mice (**Figure 1**).

LD amastigotes are lipid auxtrophs [11, 14], and this severe dyslipidemia in response to LD-R-infection can be logically interpreted in terms of LD-R’s increase need to support their aggravated intracellular proliferation in the infected-host. In liver-resident macrophages (Kupffer cells), primary sites for LD infection and host cholesterol synthesis, we observed significantly higher number of LD-R amastigotes and parasitophorous vacuoles formation (PV), compared to LD-S-infection, indicating LD-R’s better capacity to utilize host cholesterol (**Figure 1, S1**). Also, in hyperlipidemic Apoe^-/-^ mice, LD-R-infection led to organ parasite overload compared to LD-S-infection, confirming LD-R’s increased dependence on extracellular lipid sources to sustain its aggressive proliferation (**Figure 2**). Thus, next we focused on investigating LD-R’s ability to differentially regulate host’s immune-metabolic status. Previous reports suggested that LD with altered drug-responsiveness can modulate their host differentially resulting in different disease outcome [3, 61]. Our results confirmed that while both LD-R and LD-S promastigotes initially quench cholesterol from the host plasma membrane similarly (**Figure S2**), LD-R amastigotes accumulate significantly more cholesterol and fatty acids in their PV by specifically promoting increased endocytosis of LDL as the infection progresses (**Figure 3, S3**). This possibly also explains the similar initial attachment and infectivity of LD-S and LD-R promastigotes with the host MFs again indicating towards their ability to infect their host equally (**Figure 1, S1, Movie 1**).

Interestingly, this LD-R-induced LDL-endocytosis is independent of the host’s LDL receptor and relies on fluid-phase endocytosis, involving significant actin cytoskeleton reorientation in LD-R-infected-MFs (**Figure 4, S4**). It is worth mentioning that LD-R-infection has been reported to induce high levels of IL-10 [3], which is linked with enhanced fluid-phase endocytosis [42].

Importantly, we also noted an increased convergence of endocytosed LDL-vesicles with lysosomal associated marker protein (LAMP-1), along with higher LAMP-1 expression around LD-R amastigotes as compared to LD-S ones (**Figure 5**). Lysosomal acid hydrolases metabolize LDL into bioavailable cholesterol, and higher fusion of lysosomal vesicles with LD-R-PV explains the significant Filipin-cholesterol staining around LD-R amastigotes (**Figure 5**). These findings not only confirm that endocytosed-LDL can be better utilized by LD-R amastigotes but probably also indicates towards a rapid maturation of LD-R-PV due to increased LDL-vesicle fusion [13, 36].

Although available RNAseq analysis (**Figure 6**) did not support this increased expression of l*amp-1* in the transcript level, it did reflect a notable upregulation of vesicular fusion protein (VSP) *vamp8* and *stx1a* in response to LD-R-infection. How, LD infection can regulate LAMP-1 expression, and the role of VSPs in LDL-vesicle fusion with LD-R-PV is worthy of further investigation. It is possible and has been earlier reported that LD infection can regulate host proteins expression through post transcriptional and post translational modifications [62–64]. It is tempting to speculate that LD-R amastigote might be promoting an increased lysosomal biogenesis through any such mechanism to increase supply of bioavailable cholesterol through action of lysosomal acid hydrolases on LDL. Additionally, RNAseq analysis also showed suppressed expression of *npc-1* in LD-R-infected-MFs. NPC-1 is crucial for mobilizing vesicular cholesterol to the cytoplasm and plasma membrane [44], and its suppressed expression correlates with higher cholesterol availability for aggressive LD-R proliferation (**Figure 6**).

Besides LDL endocytosis, *de novo* cholesterol synthesis could also serve as a lipid source for intracellular LD amastigotes [65]. Interestingly, LD-R-infected macrophages exhibited suppressed expression of HMGCR and SREBP2, the key regulators of de novo cholesterol biosynthesis. Additionally, SREBP2 showed minimal nuclear translocation in LD-R-infected macrophages compared to LD-S-infected cells, (**Figure 6**), consistent with reports of a shutdown in de novo cholesterol synthesis in MFs with high endocytosed LDL accumulation [66]. Notably, SREBP2 regulates NPC-1 expression [46] and has been and has been shown to get activated in response to LD-S infection [66]. Further RNA sequencing data also revealed a significant downregulation of *hmgcs* (3-hydroxy-3-methylglutaryl-CoA synthase) in LD-R infected PECs as compared to LD-S infection. Downregulation of HMGCS which catalyzes the condensation of acetyl-CoA with acetoacetyl-CoA to form 3-hydroxy-3-methylglutaryl-CoA (HMG-CoA), which serves as an intermediate in both cholesterol biosynthesis and ketogenesis further supports our observation that LD-R-infected PECs preferentially rely on endocytosed low-density lipoprotein (LDL)-derived cholesterol rather than *de novo* synthesized cholesterol to support their metabolic needs.

Surprisingly, LD-R-Infected-MFs showed minimal inflammatory response despite high lipid accumulation, due to their ability to successfully exclude uptake of oxidized LDL (ox-LDL). Low expression of MSR-1, a scavenger receptor of ox-LDL, in the liver of LD-R-infected mice explained this observation, leading to the accumulation of neutral lipid droplets around LD-R amastigotes (**Figure 7, S5, Movie 4**). Expansion Microscopy (ExM) [56] along with TEM confirmed significant assimilation of these neutral lipids into LD-R amastigotes (**Figure 8**). Several intracellular pathogens like *Mycobacterium*, *Toxoplasma*, HCV have been reported to transfer stored triacylglycerols and sterol esters from lipid droplets to support their proliferation [10, 67, 68]. While the understanding of lipid uptake and metabolizing genes in LD is limited, a recent genome-wide CRISPR screen in *L. Mexicana* [69], have identified such potential candidates “essential” for parasite survival. Close proximity lipid droplets with LD-R amastigotes makes it tempting to speculate existence of lipid droplets contact sites [13], or possible copy number variations in these lipid metabolizing genes [70], providing survival advantages to LD-R. This speculation is further supported by the fact that although clinical LD-R isolates with neutral lipid droplet accumulation in infected MFs displayed increased unresponsiveness to Amphotericin-B (Amp-B), lab-generated Amp-B unresponsive promastigotes failed to do so in infected MFs (**Figure 8**). Fact that LD-R amastigotes regained their responsiveness towards Amp-B in low lipid medium or in presence of Aspirin [53], further confirms that eliminating lipid droplets around LD-R amastigotes might be a weapon of choice to combat treatment failure against Amp-B [27, 28, 71]. Our results thus suggest that natural clinical LD-R-isolates have developed an inherent adaptability to gradually develop cross-resistance towards Amp-B, probably as an additional consequence of their ability to accumulate increased host derived lipids as a rapidly proliferating intracellular amastigote (represented graphically in **Figure 9**). Our work explains increasing Amp-B treatment failure among VL patients [27, 28, 71] and correlates with low serum LDL and cholesterol levels in patients relapsing after Amp-B treatment (**Figure 9, Table 2**).

**Figure 9:**
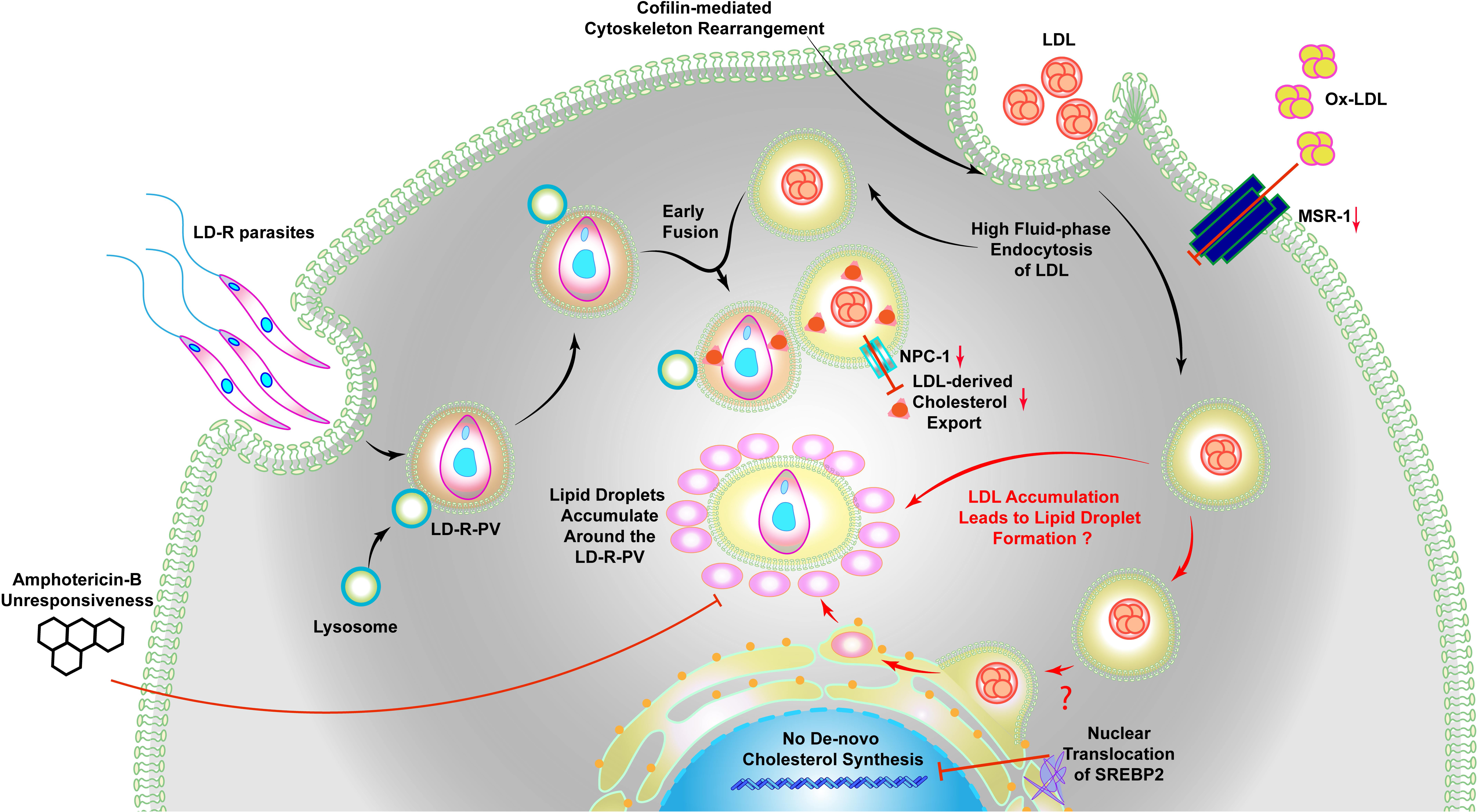
Scrutinized uptake of host lipids results in breakdown of Amphotericin-B responsiveness in clinical LD isolates with primary resistance to antimony. LD-R reside within membrane-bound parasitophorous vacuoles (PVs) and actively modulate host actin cytoskeleton dynamics through cofilin-mediated depolymerization, inducing high fluid-phase endocytosis of LDL. The internalized LDL-containing vesicles subsequently fuse with LD-R-associated PVs, facilitating increased lipid acquisition essential for their aggressive intracellular proliferation. In these fused membranes, the vesicular cholesterol export protein NPC-1 is downregulated, effectively sequestering LDL-derived cholesterol within the PV and preventing its release into the host cytoplasm. Concurrently, LDL-derived cholesterol esters either directly or getting processed in the endoplasmic reticulum (ER) can lead to the formation of lipid droplets (as presented by **?,** since it needs further investigation. Lipid droplets then accumulate in and around the LD-R-PV. This lipid droplet accumulation is a key factor in rendering LD-R infections unresponsive to Amphotericin-B. Additionally, LD-R regulate inflammatory responses linked to oxidized lipid (Ox-LDL) accumulation by downregulating macrophage scavenger receptor 1 (MSR-1). The elevated LDL influx also leads to a shutdown of *de novo cholesterol* biosynthesis by restricting the nuclear translocation of sterol regulatory element-binding protein 2 (SREBP2).

## Methods

### Reagents

Laurdan (Merck), photoRED-LDL (Thermo Fisher Scientific), NBD Cholesterol (Sigma), Nile red (Sigma-Aldrich), Oil red O (Thermo Fisher Scientific), Filipin (Sigma-Aldrich), Collagenase (Gibco) were purchased. Tissue culture plasticware were purchased from NUNC (Roskilde, Denmark). Invitrogen Amplex-Red assay kit was purchased from Thermo Fisher Scientific. Primary antibodies required for the experiments were anti-ATP6V0D2 (ABclonal), anti-MSR1 (ABclonal), anti-LAMP1 (Thermo Fisher Scientific), anti-SREBP2 (Thermo Fisher Scientific), anti-NPC1 (Santa Cruz Biotechnology), anti-HMGCR(Santa Cruz Biotechnology), anti-LDLr (Santa Cruz Biotechnology), anti-KMP-11(Invitrogen), goat anti-mouse IgG Alexa Fluor 488 (Thermo Fisher Scientific), goat anti-mouse IgG Alexa Fluor 594 (Thermo Fisher Scientific), goat anti-Rabbit IgG-HRP (Thermo Fisher Scientific), goat anti-Mouse IgG-HRP (Thermo Fisher Scientific), β actin (Cell Signalling Technology), CD11b (Cell Signalling Technology), Coffilin (Cell Signalling Technology), and phospho-Coffilin (Cell Signalling Technology). RhodaminePhalloidin reagent was purchased from Thermo Fisher Scientific and ELISA Kits for IFN-**γ** were purchased from R&D Systems, Aspirin generic composition, SAG (Albert David)

### Animals

BALB/c, C57BL/6 and *Apoe^-/-^* mice were maintained and bred under pathogen-free conditions with food supplements and filtered water. All experiments were conducted in accordance to Institutional Animal Ethics Committee (IAEC), IIT Kharagpur guidelines India (IAEC/BI/1 55/2021) and approved by National Regulatory Guidelines issued by Committee for the Purpose of Supervision of Experiments on Animals, Ministry of Environment and Forest, Government of India.

### Ethical statement

In this study, all procedures involving human participants were conducted in accordance with the ethical standards of the 1964 Helsinki Declaration and approval of the “Institutional Human Ethical Committee” of ICMR-Rajendra Memorial Research Institute of Medical Sciences, Patna, India (Approval No: RMRI/EC/20/2020). Following information about the potential risks, benefits, and the investigational nature of the study consent from all study participants was obtained in the informed consent form.

### Parasite Cultures and Maintenance

*Leishmania donovani* (LD) isolates used in this study are natural clinical isolates obtained from the patients MHOM/IN/09/BHU569/0, MHOM/IN/2009/BHU575/0, MHOM/IN/10/BHU814/1, MHOM/IN/83/AG83, MHOM/IN/09/BHU777/0, MHOM/IN/80/DD8 [2, 3], and were maintained in BALB/c mice as per institute biosafety guidelines (BT/BS/17/27/97-PID). Susceptibility of each of these LD isolates against antimony were checked and confirmed against previously reported EC50 value for antimony before they were included in the study (This can be reported in a common table, Table 1). Lab generated Amp-B resistant strain (DD8-AmpB) strain was kind from Dr. Arun Kumar Haldar (CSIR-CDRI). Amastigotes were isolated by macerating the spleen of 21 days infected mice under sterile conditions, and subsequently transformed into promastigotes by maintaining in M199 (Gibco) supplemented with 5% (v/v) heat-inactivated fetal bovine serum (FBS) (Gibco) and 1% (v/v) Penicillin Streptomycin solution (Gibco) under drug selection.

**Table-1:** EC50 against SAG (Antimonial) for the LD strains used in this study.

| Strain Name | EC50 value for<br>SAG($\mu\text{g/ml}$) in PEC | EC50 value for<br>SAG($\mu\text{g/ml}$) in KC |
| --- | --- | --- |
| MHOM/IN/2009/BHU575/0 | 22.73±3.7 | 23.89±2.1 |
| MHOM/IN/10/BHU814/1 | 30.3±7.2 | 28.5±5.3 |
| MHOM/IN/83/AG83 | 1.52±1.2 | 3.3±2.2 |
| MHOM/IN/80/DD8 | 2.1±1.2 | 3.2±1.9 |
| DD8-AmpB | 1.8±2.1 | 2.2±1.3 |
| BHU575-EGFP | 21.45±3.2 | 22.81±3.1 |
| AG83-EGFP | 2.1±1.4 | 2.4±3.1 |

**Table-2:** Lipid profile of VL patients.

| Serum Code | Age | Sex | Time of Blood Collect | Infection status | Treatment received | Patient status | LDL (mg/dl) | HDL (mg/dl) | Cholesterol (mg/dl) | Triglycerides (mg/dl) |
| --- | --- | --- | --- | --- | --- | --- | --- | --- | --- | --- |
| RMRI N1 | 45 | F | Before | Control population from endemi | NA | NA | 45.3501585 | 46.6365931 | 101.6949153 | 177.258567 |
| RMRI N6 | 70 | F | Before | Control population from endemi | NA | NA | 110.855478 | 50.3762712 | 111.8644068 | 206.5420561 |
| RMRI N7 | 45 | F | Before | Control population from endemi | NA | NA | 75.5620787 | 41.8711864 | 84.40677966 | 165.4205607 |
| RMRI N8 | 19 | M | Before | Control population from endemi | NA | NA | 83.946038 | 62.9533123 | 100.3389831 | 224.9221184 |
| RMRI N11 | 37 | M | Before | Control population from endemi | NA | NA | 72.4549921 | 120.914196 | 134.2372881 | 244.5482866 |
| RMRI N12 | 60 | M | Before | Control population from endemi | NA | NA | 52.6307448 | 113.851735 | 92.54237288 | 180.6853583 |
| RMRI N13 | 30 | M | Before | Control population from endemi | NA | NA | 66.7533281 | 48.3413249 | 138.3050847 | 236.4485981 |
| RMRI N14 | 22 | M | Before | Control population from endemi | NA | NA | 47.8939778 | 49.6807571 | 100 | 260.7476636 |
| RMRI N15 | 37 | F | Before | Control population from endemi | NA | NA | 46.4904913 | 45.9059937 | 108.8135593 | 219.6261682 |
| RMRI C1 | 20 | F | Before | Confirmed VL, Hospitalization | Ambisome Single dose | Cured | 110.389045 | 44.2700565 | 112.5423729 | 200.623053 |
| RMRI C2 | 62 | F | Before | Confirmed VL, Hospitalization | Ambisome Single dose | Cured | 121.738904 | 64.5514124 | 145.4237288 | 301.8691589 |
| RMRI C3 | 15 | M | Before | Confirmed VL, Hospitalization | Ambisome Single dose | Cured | 76.1839888 | 36.7463277 | 95.59322034 | 185.6697819 |
| RMRI C4 | 31 | M | Before | Confirmed VL, Hospitalization | Ambisome Single dose | Cured | 72.141573 | 41.1079096 | 116.9491525 | 241.7445483 |
| RMRI C5 | 52 | M | Before | Confirmed VL, Hospitalization | Ambisome Single dose | Cured | 98.1063202 | 48.1954802 | 117.2881356 | 282.2429907 |
| RMRI C6 | 18 | F | Before | Confirmed VL, Hospitalization | Ambisome Single dose | Cured | 68.8765449 | 36.7463277 | 96.61016949 | 200.623053 |
| RMRI C7 | 45 | F | Before | Confirmed VL, Hospitalization | Ambisome Single dose | Cured | 110.544522 | 55.6101695 | 124.4067797 | 381.9314642 |
| RMRI C8 | 32 | F | Before | Confirmed VL, Hospitalization | Ambisome Single dose | Cured | 94.8412921 | 43.0706215 | 106.1016949 | 232.3987539 |
| RMRI C18 | 66 | M | Before | Confirmed VL, Hospitalization | Ambisome Single dose | Cured | 61.4136236 | 35.9830508 | 91.86440678 | 203.4267913 |
| RMRI R14 | 8 | M | Before | Confirmed VL, Hospitalization | Ambisome Single dose | Relapse | 79.2935393 | 83.6333333 | 96.94915254 | 262.9283489 |
| RMRI R16 | 49 | M | Before | Confirmed VL, Hospitalization | Ambisome Single dose | Relapse | 56.9047753 | 102.279096 | 63.05084746 | 209.0342679 |
| RMRI R17 | 18 | M | Before | Confirmed VL, Hospitalization | Ambisome Single dose | Relapse | 59.7033708 | 101.079661 | 83.38983051 | 217.1339564 |
| RMRI R18 | 59 | M | Before | Confirmed VL, Hospitalization | Ambisome Single dose | Relapse | 63.4348315 | 94.319209 | 84.40677966 | 304.6728972 |

### Cell Culture and Infection

Peritoneal exudate cells (PECs) were harvested from BALB/c mice as mentioned in already published protocol, with slight modifications [72]. After 48Hrs of 4% starch intraperitoneal injection, PECs were isolated and plated on 35-mm tissue culture petri dishes, sterile 22-mm square coverslips in 35-mm petri dishes or 24-well plates at a density of 1×10^6^, 1.5×10^5^ or 1.5×10^5^, respectively, all in RPMI 1640 medium (Gibco) supplemented with 10% (v/v) heat-inactivated FBS, 1% (v/v) Penicillin Streptomycin solution. The cells were allowed to adhere for 48Hrs at 37°C in 5% CO_2_ before the infection process.

### *In vivo* infection

For *in vivo* infections, healthy female C57BL/6 mice or *Apoe^-/-^*mice (4-6 weeks old and 20-25 g each) were injected via tail vein route with 1×10^7^ GFP-expressing *L. donovani* stationary phase metacyclic promastigotes (suspended in 100 μl 1X PBS) as reported previously [73]. After infection mice were kept in atherogenic diet till sacrifice. Each group contained five mice that were sacrificed 11 days’ post-infection (p.i). Splenocytes were isolated and GFP positive population was measured through flow cytometry. Each experiment was performed in triplicate.

### Isolation of Kupffer cells

Kupffer cells (KCs) were harvested from mice livers following a previously published protocol, with slight modifications [74]. Wild-type BALB/c mice were anesthetized, and warm PBS containing 0.05% collagenase IV was used for liver perfusion. The livers were swiftly excised and minced into small fragments on an ice bath. Tissue digestion was performed using a 0.1% Type IV Collagenase solution in RPMI 1640 for 30 minutes at 37°C with intermittent shaking for every 10 minutes. Following digestion, KCs were separated from hepatocytes and other cellular components through sequential centrifugation steps and plated on well plates, as required, for 4Hrs in RPMI 1640 media supplemented with 10% (v/v) Penicillin Streptomycin and 10% (v/v) FBS. After 24Hrs of resting at 37°C in 5% CO_2_, the infection studies commenced. To assess the purity of the isolated KCs by flow cytometry, KCs were plated on a 6 well plate. After 4Hrs, KCs were scraped out, centrifuged at 300g for 5 minutes and stained with monoclonal anti-F4/80 PE conjugated antibody (BD Biosciences). The KCs were then washed and fixed with 2% paraformaldehyde for 5 minutes, analyzed using a flow cytometer (ARIA FACS DIVA), and further processed using FLOW JO software for data analysis. Purified KCs were infected with stationary phase sorted metacyclic LD promastigotes as mentioned previously. PECs and KCs are collectively designated as MFs (macrophages).

### Sorting of Metacyclic LD and infection in MΦs

Metacyclic parasites were sorted through Beckman Coulter CytoFLEX cell sorter, as per manufacturer’s protocol. The LD promastigotes were washed and resuspended in PBS before being subjected to light scatter analysis using flow cytometry. Sorted population were checked with forward scatter (FSC) versus side scatter (SSC) and dot plot was generated to represent the acquisition of 10,000 events using a FACS Aria II system. Required number of sorted metacyclic LD were used to perform subsequent MF and mice infections. MFs were infected with stationary-phase Flow sorted metacyclic LD promastigotes at a MOI 1:10 ratio (in some cases 1:5 and 1:20) for 4Hrs which was considered as the 0^th^ point of the infection. MFs was then washed to remove any extracellular loosely attached parasites and incubated further as per experimental requirement.

### Intracellular amastigote load determination

To enumerate intracellular amastigote load, resting stage PECs and KCs on glass coverslips were infected with sorted metacyclic LD promastigotes of various strains at a ratio of 1:10, as described in previous protocols [3, 75]. Following a 4Hrs incubation, the PECs or KCs were washed and subsequently cultured for specific durations. Coverslips were retrieved after specific time points, rinsed with PBS, fixed using methanol, and stained with Giemsa. These prepared coverslips were then affixed onto glass slides and examined utilizing a light microscope (Olympus). Parasite quantification was conducted by counting amastigotes per 100 macrophages. Image processing related to Figure 1A and Figure S1 D was performed using Fiji by separating the spectral overlap between the light and dark stained region to Giemsa stained MFs. Next, for better visualization, pseudo colours were assigned to represent nucleus (white) and cytoplasm (red) with a black background. Colour separated images were then merged showing LD within PV.

### Raman Spectroscopy of LD-infected-KCs

KCs were plated on Raman-grade stainless steel plate and kept for 24Hrs in a humidified environment at 37°C with 5% CO_2_, followed by infection with LD as already described. At 24Hrs p.i., the KCs were washed using PBS, fixed with 2% paraformaldehyde and proceeded to perform Raman spectroscopy (Witec, alpha300R). The Raman grade stainless steel plates were positioned on the microscope stage and focused with a 100× air objective and measurements were performed using a 532 nm laser source producing a laser spot of approximately 0.3 µm with 20mW laser power. Spectral measurements were obtained through the Project FIVE interface (Witec, Germany) after setting the grating to 600 gratings/mm. The spectral resolution was set to 2.87 cm^−1^. The Raman spectra were taken with integration time set to 5 seconds, and ten accumulations were selected after confirming its effectiveness on the KCs being studied. Baseline correction and other spectral denoising procedures were carried out using MATLAB 2017b (MathWorks), and the processed spectra were analysed as described preciously [37, 39]. For confocal Raman spectroscopy, spectral data were acquired from individual cells at 1000× magnification using a 100 × 100 μm scanning area, following previously established specifications. After spectral acquisition, distinct Raman shifts corresponding to specific biomolecular signatures were extracted for further analysis. These included: Cholesterol (535–545 cm⁻¹), Nuclear components (780–790 cm⁻¹), Lipid structures (1262–1272 cm⁻¹), Fatty acids (1436–1446 cm⁻¹) Following spectral extraction, pseudo-color mapping was applied to highlight the spatial distribution of each biomolecular component within the cell. These processed spectral images are presented in Figure 3D1, where the first four panels illustrate the individual biomolecular distributions. A merged composite image was then generated to visualize the co-localization of these biomolecules within the cellular microenvironment, with the final panel specifically representing the spatial distribution of key biomolecules.

### Quantification of membrane cholesterol with Amplex Red assay kit

The KC membrane preparation was conducted following the method described previously [20], with slight modifications. KCs were ruptured by repeated freezing and thawing cycles followed by probe sonicating for 4 cycles with 30secs burst and 30sec gap. The resulting homogenate was then centrifuged at 900g for 10mins at 4°C. The supernatant was filtered through a nylon mesh (100 μm), while the pellet was discarded. The filtered supernatant was centrifuged at 100,000g for 1hr at 4°C, and resulting pellet, representing native membrane, was suspended in a buffer containing 50 mM Tris (Ph 7.4), 1 mM EDTA, 0.24 mM PMSF, and 10 mM iodoacetamide. The protein content of the membrane was quantified using Bradford reagent (Biorad), following manufacturer’s protocol. The total cholesterol content was determined using an Amplex Red reagent kit [76], with the results expressed in moles of cholesterol per gram of protein.

### Parasitophorous vacuole (PV) Isolation and cholesterol measurement

Parasitophorous vacuoles (PV) were isolated using a previously outlined protocol with slight modifications [77]. 10^7^ KCs were seeded in a 100 mm plate and allowed to adhere for 24Hrs. Following this infection was performed with Leishmania donovani (LD) for 24Hrs, the infected KCs were then harvested by gentle scraping and lysed through five successive passages through an insulin needle to ensure membrane disruption while preserving organelle integrity. The lysate was centrifuged at 200 × g for 10mins at 4°C to remove intact cells and large debris. The resulting supernatant was carefully collected and subjected to a discontinuous sucrose density gradient (60%, 40%, and 20%). The gradient was centrifuged at 700 × g for 25mins at 4°C to facilitate organelle separation. The interphase between the 40% and 60% sucrose layers, enriched with PVs, was carefully collected and subjected to a final centrifugation step at 12,000 × g for 25mins at 4°C. The supernatant was discarded, and the resulting pellet was enriched for purified parasitophorous vacuoles, suitable for downstream biochemical and molecular analyses. Cholesterol and protein contents in PV were determined by an Amplex Red assay kit and Bradford assay, respectively. Resulting data were represented as micrograms of cholesterol per microgram of protein.

### GC-MS analysis of LD-S and LD-R-PV

Following a 24Hrs infection period, KCs were harvested, washed with phosphate-buffered saline (PBS), and pelleted. Subsequent to this, PV isolation was carried out using the previously described protocol [36]. After PV isolation Bradford assay was carried out for normalizing the protein concentration. The resulting equal volume of PV pellet was suspended in 20 ml of dichloromethane: methanol (2:1, vol/vol) and incubated at 4°C for 24hours. After centrifugation (11,000 g, 1 hour, 4°C), the supernatant was checked through thin layer chromatography (TLC) and subsequently evaporated under vacuum. The residue and pellet were saponified with 30% potassium hydroxide (KOH) in methanol at 80°C for 2 hours. Sterols were extracted with n-hexane, evaporated, and dissolved in dichloromethane. A portion of the clear yellow sterol solution was treated with N, O-bis(trimethylsilyl)trifluoroacetamide (BSTFA) and heated at 80°C for 1 hour to form trimethylsilyl (TMS) ethers. Gas chromatography/mass spectrometry (GC/MS) analysis was performed using a Varian model 3400 chromatograph equipped with DB5 columns (methyl-phenylsiloxane ratio, 95/5; dimensions, 30 m by 0.25 mm). Helium was used as the gas carrier (1 ml/min). The column temperature was maintained at 270°C, with the injector and detector set at 300°C. A linear gradient from 150 to 180°C at 10°C/min was used for methyl esters, with MS conditions set at 280°C, 70 eV, and 2.2 kV[78].

### Cytokine measurement

Supernatants from KCs infected with stationary-phase LD co-culture with T cells, isolated from the mice infected with LD. For detection of IFN-**γ** from Apoe-/-, splenocytes isolated from LD infected mice and cultured with SLA. IFN-**γ** levels in the supernatants were measured using a sandwich ELISA Kit (R&D Systems) as per manufacturer’s protocol. Detection limit for these kits were 8pg/ml for IFN-**γ**. Cytokine levels were determined by measuring the OD at 450 nm using a MultiSkan FC microplate photometer (Thermo Fisher Scientific). Data was demonstrated as mean ± SD of all five individual experiments.

### Western Blot Analysis

Lysates of LD-infected PECs and KCs were prepared, and western blotting was performed for different proteins with endogenous control β-actin (Cell Signaling Technology). Blots were probed with specific antibodies. Binding of secondary HRP-labeled goat anti-rabbit or goat anti-mouse Abs was analyzed using SuperSignalR West Pico or West Dura Chemiluminescent substrate (Pierce).

### Confocal and Structural Illumination Microscopy (SIM)

PECs and KCs were cultured on 13-mm diameter glass coverslips at a density of 2*10^5^, and infected with LD as mentioned previously. After varying time intervals, infected PECs or KCs were rinsed with PBS, fixed with 2% paraformaldehyde, and permeabilized using 0.2% Triton/PBS for 20mins on an orbital shaker. Non-specific binding was blocked with 2% BSA in 0.2% Triton/PBS for another 20mins on an orbital shaker. The PECs and KCs were then incubated with primary antibodies for 1Hr, followed by washing and staining with IgG Alexa Fluor 594 or IgG Alexa Fluor 488, as required. Mounting was performed using Fluromount-G-DAPI (Thermo Scientific). Images were captured using Olympus confocal microscopy (FV3000) and super-resolution microscopy ZEISS (SIM) with a 63× objective. 3D image visualization processed through Fiji 3D viewer and 3D project plugin. Live cell microscopy was performed using ZEISS phase contrast microscope and ZEISS Apotome. MFs were plated on confocal dish and incubated with photoRED-LDL for live cell imaging of LDL uptake (**Movie 3)**.

### Image processing and analysis

Image processing and analysis were conducted using Fiji (ImageJ). For optimal visualization, Giemsa-stained macrophages (MΦs) were represented in grayscale to enhance contrast and structural clarity. To improve the distinction of different fluorescent signals, pseudo-colors were assigned to fluorescence images, ensuring better differentiation between various cellular components. For colocalization analysis (Figures 3, Figure 5, Figure 6, and Figure S2), we utilized the RGB profile plot plugin in ImageJ, which allows for the precise assessment of signal overlap by generating fluorescence intensity profiles across selected regions of interest. This approach provided quantitative insights into the spatial relationship between labelled molecules within infected cells. Additionally, for analysing the distribution of cofilin in Figure 4, the ImageJ surface plot plugin was employed. This tool enabled three-dimensional visualization of fluorescence intensity variations, facilitating a more detailed examination of cofilin localization and its potential reorganization in response to infection.

### Lipid droplet staining

LD-infected-KCs were first fixed using 2% paraformaldehyde (PFA) and subsequently permeabilized with 0.25% Triton X-100. Coverslips were then treated with Nile red (Sigma, diluted 1:100) in PBS for at least 1Hr to stain lipids, followed by DNA staining with DAPI. The coverslips were mounted onto glass slides using Fluoromount-G and visualized with confocal microscope (LEICA STELLARIS 5). 3D stacks were acquired to visualize distribution of lipid droplets. Images were processed and analysed, and stacks were merged to reconstruct 3D images using Fiji.

### Ultrastructure expansion microscopy

Ultrastructure Expansion Microscopy (U-ExM) was conducted on LD-infected-MΦs following established experimental protocols [56]. Protein crosslinking was initiated by immersing the coverslip in a solution containing 1.4% formaldehyde, 2% acrylamide, and PBS for 5Hrs at 37°C. Monomer solution (19% sodium acrylate, 10% acrylamide, and 0.1% N,N’-methylenbisacrylamide in PBS) was mixed with 10% TEMED and 10% APS solution on ice and added to 6-well pates. Gelation was performed on ice, with cell-plated coverslip placed face down on the gelation solution in a humid chamber for 5 minutes at 37°C for 1 hour. The gel and coverslip were together transferred to a 6-well plate filled with denaturation buffer (200 mM SDS, 200 mM NaCl, and 50 mM Tris pH 9.0) and incubated face up for 15mins at room temperature under agitation, followed by incubation at 95°C for 1.5Hrs. The first expansion phase involved incubating the gel three times in ddH2O for 30mins each. Previously mentioned IFA steps were adopted for antibody staining. Images were captured using an Olympus confocal microscope (FV3000) equipped with a 63X 1.4 NA oil objective and Small Volume Computational Clearing mode was used to obtain deconvolved images. 3D stacks were acquired and analyzed using Fiji software.

### Transmission electron microscopy (TEM)

For transmission electron microscopy, PECs infected with the LD-R strain were fixed after 24 hours of infection using 2.5% glutaraldehyde, an electron microscopy (EM) grade fixative. The cells were then processed according to previously established protocols[58]. Following this, the samples were examined using a JEM-F200 transmission electron microscope, allowing for lipid droplet imaging and detailed visualization of the cellular structures.

### *In-vitro* cholesterol trafficking assay

KCs was first treated with NBD-cholesterol for 16Hrs, allowing the fluorescent cholesterol analog to integrate into their membranes. Next, KCs were washed with RPMI 1640 and cultured for again 2Hrs in the same medium to eliminate any residual fluorescence from the background. These labelled KCs were infected with LD for 4Hrs and washed to remove loosely attached MFs. The KCs were then either immediately fixed with 2% paraformaldehyde (PFA) or left untreated to serve as a control group. Finally, the fixed MFs were examined under a confocal microscope.

### Flow cytometry to determine NBD-cholesterol

KCs were first treated with NBD-cholesterol as previously described. LD parasites were isolated after 2Hrs of attachment to the KCs. The quenching of NBD-cholesterol was measured by flow cytometric analysis, using an unstained control.

### Total internal reflection fluorescence (TIRF) microscopy

To visualize membrane fluidity, TIRF microscopy was performed [79]. Briefly, KCs were plated on coverslips, and after 24Hrs of LD-infection, KCs were fixed with 2% PFA. KCs were stained with Laurdan dye and visualized with Olympus cell TIRF microscopic system. Representative images of three independent biological experiments have been provided.

### RNAseq analysis

RNA-Sequencing was performed utilizing external service by Bionivid Project for LD-S and LD-R-infected-PECs, following 24Hrs of infection keeping uninfected MΦs as control (Bioproject. Average expression of differentially expressed genes related to lipid metabolism between LD-S and LD-R-infected-KC is represented as heat map. RAW sequencing reads have been deposited to Indian Nucleotide Data Archive (https://inda.rcb.ac.in/home) with accession number INRP000146.

### Lipid profiling from serum

Lipid profile analysis of patient serum was conducted using lipid estimation kits employing the enzymatic method for the determination of cholesterol, triglycerides, HDL, and LDL. For BALB/c mice, blood was collected following anesthesia and prior to euthanasia. The blood samples were maintained at 4°C to allow clotting, after which the serum was isolated.

### Evaluation of EC50

KCs infected with LD parasites were treated with SAG or Amp-B 24Hrs post-infection and incubated for an additional 24Hrs. At the experimental endpoints, coverslips were washed with PBS, air-dried, and fixed with 100% ice-cold methanol. Subsequently, the coverslips were stained with Giemsa solution and examined microscopically. To quantify the number of amastigotes, one hundred KCs per coverslip were counted. [2]. The average of three untreated coverslips was taken as 100% control, and the percentage inhibition of infected KCs in treated cultures was calculated. EC50 values for each isolate were estimated against each drug [2]. Results were also expressed as EC50.

### Aspirin treatment and amastigote load determination

KCs were infected with LD parasites and, after 4Hrs, treated with 5μM Aspirin for a duration of 24 hours. Following this treatment, the KCs were washed and subsequently treated with Amp-B at a specific dose for an additional 24 hours. Coverslips were then washed with PBS, air-dried, and fixed with 100% ice-cold methanol. After fixation, the coverslips were stained with Giemsa solution and examined microscopically. To quantify the number of amastigotes, one hundred KCs per coverslip were counted. [2]

### siRNA mediated Knock down (KD)

For all siRNA transfections, Lipofectamine® RNAiMAX Reagent (Life Technologies, 13778100) specifically designed for knockdown assays in primary cells was used according to the manufacturer’s instructions with slight modifications. PECs were seeded into 24-well plates at a density of 1×10^5^ per well, and incubated at 37°C with 5% CO2. The transfection complex, comprising (1µl Lipofectamine® RNAiMAX and 50µl Opti MEM) and (1 µl siRNA and 50µl Opti MEM) mixed together directly added to the incubated PECs. Gene silencing was checked by IFA and by Western blot as mentioned previously.

### Blood samples collection

Blood samples were collected from a total of 22 individuals spanning a diverse age range (8 to 70 years) by RMRI, Bihar, India. Among these, nine samples were obtained from healthy individuals residing in endemic regions to serve as controls. Serum was isolated from each blood sample through centrifugation, and the lipid profile was subsequently analysed using a specialized diagnostic kit (Coral Clinical System) following the manufacturer’s protocol.

### Statistical analysis

All statistical analyses were performed using GraphPad Prism 8 on raw datasets to ensure robust and reproducible results. For datasets involving comparisons across multiple conditions, one-way or two-way analysis of variance (ANOVA) was conducted, followed by Tukey’s post hoc test to assess pairwise differences while controlling for multiple comparisons. A 95% confidence interval (CI) was applied to determine the statistical reliability of the observed differences. For non-parametric comparisons across multiple groups, Wilcoxon rank-sum tests were employed, maintaining a 95% confidence interval, which is particularly useful for analysing skewed data distributions. In cases where only two groups were compared, Student’s t-test was used to determine statistical significance, ensuring an accurate assessment of mean differences. All quantitative data are represented as mean ± standard error of the mean (SEM) to illustrate variability within experimental replicates. Statistical significance was determined at P ≤ 0.05. Notation for significance levels: *P ≤ 0.05; **P ≤ 0.001; ***P ≤ 0.0001.

## Acknowledgments

SP acknowledges CSIR-UGC (2020-2021) and PMRF (2021-2024) fellowship. DD is a recipient of GATE fellowship. DM is a recipient of CSIR fellowship, SC is a recipient of GATE fellowship. Authors would like to acknowledge IIR lab, SMST, IITKGP for their help in kupffer cell isolation and Flow cytometric analysis, Confocal Microscope facility under DST-FIST grant conferred on the School of Bioscience, File no. SR/FST/LS-I/2019/595(C), and Confocal facility of Department of Biotechnology, DST-FIST, Govt. of India. Authors would like to thank Prof. Syamal Roy, IICB, India for providing all the LD strains used in this study. Prof. Yasuyuki Goto, The University of Tokyo, Japan for allowing experiments related to live cell video microscopy in his lab. Authors would like to thank Dr. Praphulla Chandra Shukla for providing Apoe^-/-^ mice. Authors acknowledges CRF SMST, and CRF IITKGP. Authors would like to thanks Dr. Moumita Bhaumik, NICED, India for helping in membrane isolation experiments. Authors would like to thank all the interns of the IDI lab, SMST who help in various experiments related to this project.

## Author Contributions

SP conceptualization, experiments, analysis, MS preparation. DD performed Raman spectroscopy experiments. DM generated all the GFP, RFP expressing LD strains, SC perform experiments related to EC50, AB performed Raman spectroscopy experiments and analysis, KC did inhibitor study related to Cytochalasin-D, Latrunculin -A, RM serum collection from VL patients, AS related serum lipid profile, formal analysis of the MS, KP related to serum lipid profile, formal analysis of the MS, SD helped in formal analysis of Raman spectroscopic experiments. B.M conceptualization, analysis, conceived and directed the project and prepared the MS with input from all the authors.

## Data availability

RAW sequencing reads have been deposited to Indian Nucleotide Data Archive (https://inda.rcb.ac.in/home) with accession number INRP000146.

## Supplementary Information

**Figure S1. Kupffer cells (KC) as an *in vitro* model for LD-infection. (A)** Flow Cytometry representing % Metacyclics LD: **(i)** LD-S (AG83, DD8) and **(ii)** LD-R (BHU814, BHU575) based on FSC and SSC. Left, small box with low FSC and SSC gate for metacyclic parasites and right panel with high FSC and SSC gated procyclic parasites. **(B)** Flow cytometry of F4/80-PE positive KC from the non-parenchymal cell population isolate from murine liver. **C** Representative Giemsa-stained images LD-infected-PEC, 4 and 24Hrs p.i for **(i)** LD-S and **(ii)** LD-R strains. Giemsa Images are represented in grey scale to clearly represent LD nucleus (black arrow). Scale Bar 20μM **C (iii)** Intracellular amastigote count for 4Hrs and 24Hrs p.i. Each dot represent count from 100 infected-KC. N=6. **D** Amastigote count from Giemsa-stained image of KC-infected with LD amastigotes, isolated from the spleen of 28 days LD-infected-mice. Each dot represent mean from 100 infected-KCs. N=6. *** signifies p value < 0.0001, ** signifies p value <0.001, n.s non-significant.

**Figure S2. LD-S and LD-R quenches host membrane cholesterol during initial entry. A (i)** KCs were pre-incubated with NBD-cholesterol for 16Hrs. Subsequently, KCs were infected with RFP expressing LD-S or LD-R for 4Hrs or left uninfected and imaged. NBD-cholesterol was excited at 488nm, and LDs are presented with magenta pseudo color. White arrow indicates the NBD-cholesterol within LD while yellow arrow represents LD nucleus. Scale bar 5μm. **A (ii)** Co-localization analysis of LD (magenta), and NBD-Cholesterol (green) from A (i), marked in grey transparent box. **B (i)** Representative Flow cytometry-based analysis of NBD in LD-amastigotes isolated from LD-infected-KCs 2Hrs after infection pre-incubated with NBD cholesterol. Black Box representing % NBD positive population. **B (ii)** Graphical representation of MFI for three independent experiments as in B (i). n.s non-significant.

**Figure S3. LD-R-PV have higher accumulation of Cholesterol and Fatty acids. (A)** Detection of PV Cholesterol, isolated from LD-S or LD-R-infected KC. Data represented as μg of protein/ μg of Cholesterol, (N=6). **(B)** TLC profile of isolated lipids from 24Hrs LD-infected-PVs with Cholesterol standard kept as control. (**C)** Individual Raman spectra from six independent LD-infected-KCs **(i)** LD-S-infected and **(ii)** LD-R-infected. Lipid-related peaks are demarcated within the spectra, including those at 1.540-560 cm-1 (Cholesterol), 2. 1080-1090 cm-1 (phospholipids), 3. 1270-1280 cm-1 (triglycerides), 4. 1300-1340 cm-1 (Amide-(II) I bond), 5. 1440-1453 cm-1 (Fatty acids and Triglycerides), and 6. 1650-1660 cm-1 (Amide-I bond). **(D)** Area under the curve, and shift are calculated and plotted for each experimental condition as represented in C (i) and (i). ** signifies p value <0.001

**Figure S4. Latrunculin-A restricts extracellular LDL influx inhibiting LD-R amastigote proliferation. (A)** Western blot comparing of LDLr expression between uninfected, LD-S, and LD-R-infected-KC 24Hrs p.i. **B (i)** Expression of LDLr determined by immunofluorescence in PECs treated with LDLr siRNA and scrambled siRNA (Sc). Scale bar 10μm. **B (ii)** Western blot confirming LDLr knockdown upon SiRNA treatment. Scrambled RNA (ScRNA) was used as a negative control, while Small Interfering RNA (SiRNA) specifically targeted LDLr transcripts, TR-1 and TR-2 represent independent experimental trials. β-Actin was used as an endogenous loading control for Western blot normalization**. C** Graph representing number of LD-amastigotes in LDLr knockdown per 100 PECs for B (i) (N = 6). Data: mean ± SD. **D (i)** Cofilin and phosphorylated-Cofilin expression by Western blot in LD-infected-PEC. **D (ii)** Graphical representation of comparative expression of abundance of Cofilin and phosphorylated-Cofilin with respect to β-actin for C(i). (**E)** Giemsa-stained images of Latrunculin-A (LAT-A) and Cytochalasin-D (CYT-D) treated LD-infected-KCs in grey scale. LD nucleus represented with Black arrow and host nucleus are represented as N. Scale bar 20μm. n.s non-significant.

**Figure S5. Aspirin treatment inhibits Lipid droplets formation and increase responsiveness of LD-R amastigotes towards Amphotericin-B. (A)** Determination of lipid droplet accumulation by Nile red staining in LD-S and LD-R-infected-KC or KC left uninfected. Dotted line mark the cell periphery while dotted box cells were used for 3D reconstruction and represent in Movie 4. Scale Bar 5 μm. (**B)** Graphical representation of Lipid Droplet positive KCs in high lipid and low lipid media. Data represented as % of lipid droplet positive KCs (N=6) collected from 3 independent experiments. (**C)** Effect of Aspirin(5μM) on Lipid droplets accumulation in LD-R-infected-KCs. Independent infection was performed with LD-R^1^ and LD-R^2^ strains (N=6) and % KCs with high lipid droplets accumulation are represented. (**D)** Representative Giemsa stained images of LD-R-infected-KCs in different experimental conditions either untreated or treated with Aspirin (5μM) or Amp-B (0.36 μM) or with a combination of Aspirin (5μM) and Amp-B (0.11 μM) at 48Hrs p.i. Scale bar 20μm. *** signifies p value < 0.0001

## Movie Legends

**Movie 1:** Phase contrast video representing attached LD promastigotes to PEC 6Hrs p.i. Black arrow showing flagella of attached LD parasite. **1A** represent LD-S and **1B** represent LD-R infection. Scale bar 10μm.

**Movie 2:** Phase contrast video representing 72Hrs LD infected KCs. scale bar 20μm.

**Movie 3:** Effect of Latrunculin-A (LAT-A) on LDL uptake in LD-R infected PECs. **3A.** In absence of LAT-A. **3B.** In presence of LAT-A. LD (green), LDL (red) nucleus (blue). Scale bar 60μm

**Movie 4: 3D-**representation of lipid droplets around the LD-S amastigotes and LD-R amastigotes (S5B). Scale bar 5μm

