## Supplementary figures and images for "Scrutinized lipid utilization disrupts Amphotericin-B responsiveness in clinical isolates of *Leishmania donovani*"

### Supplementary Figures

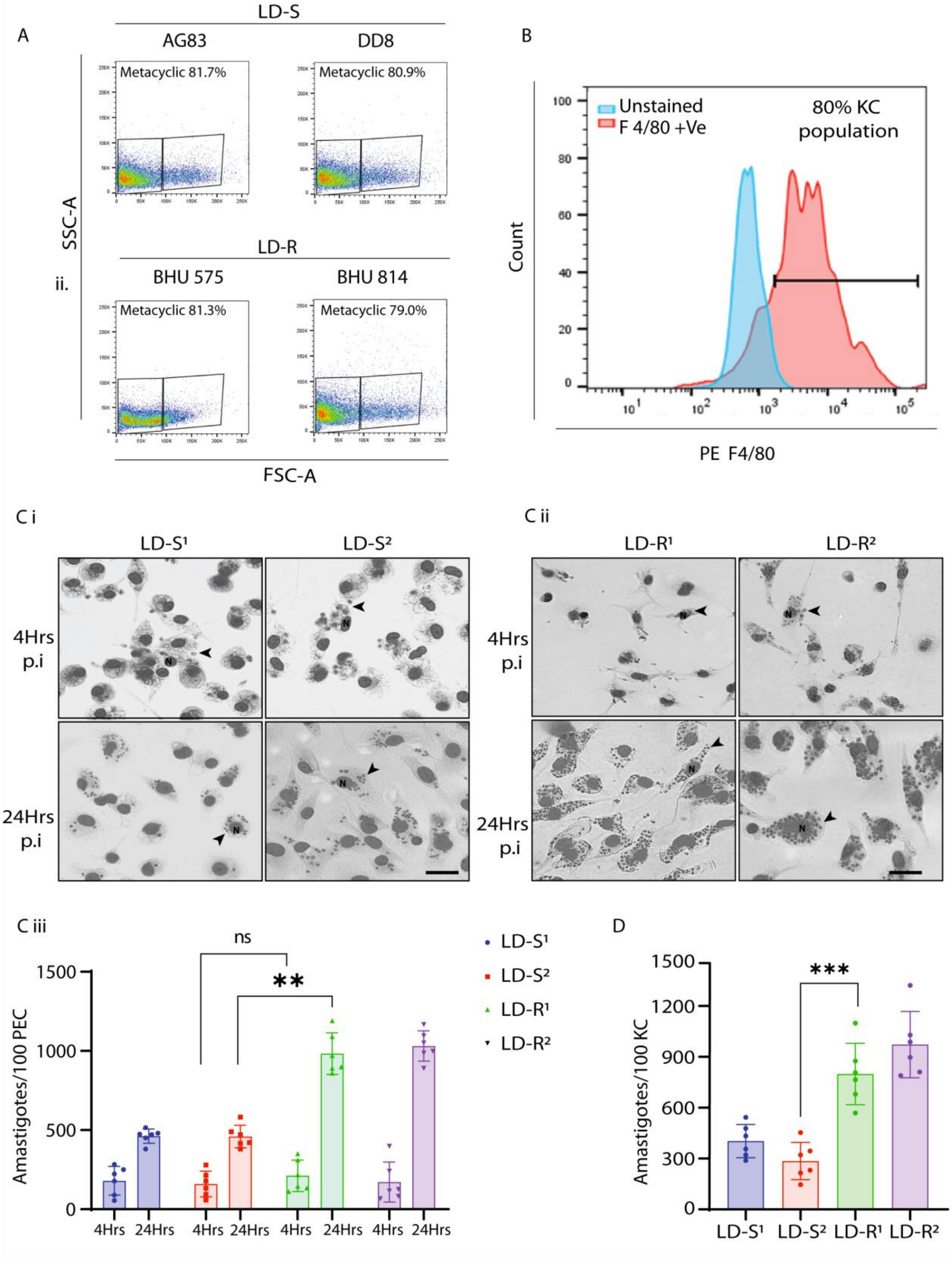

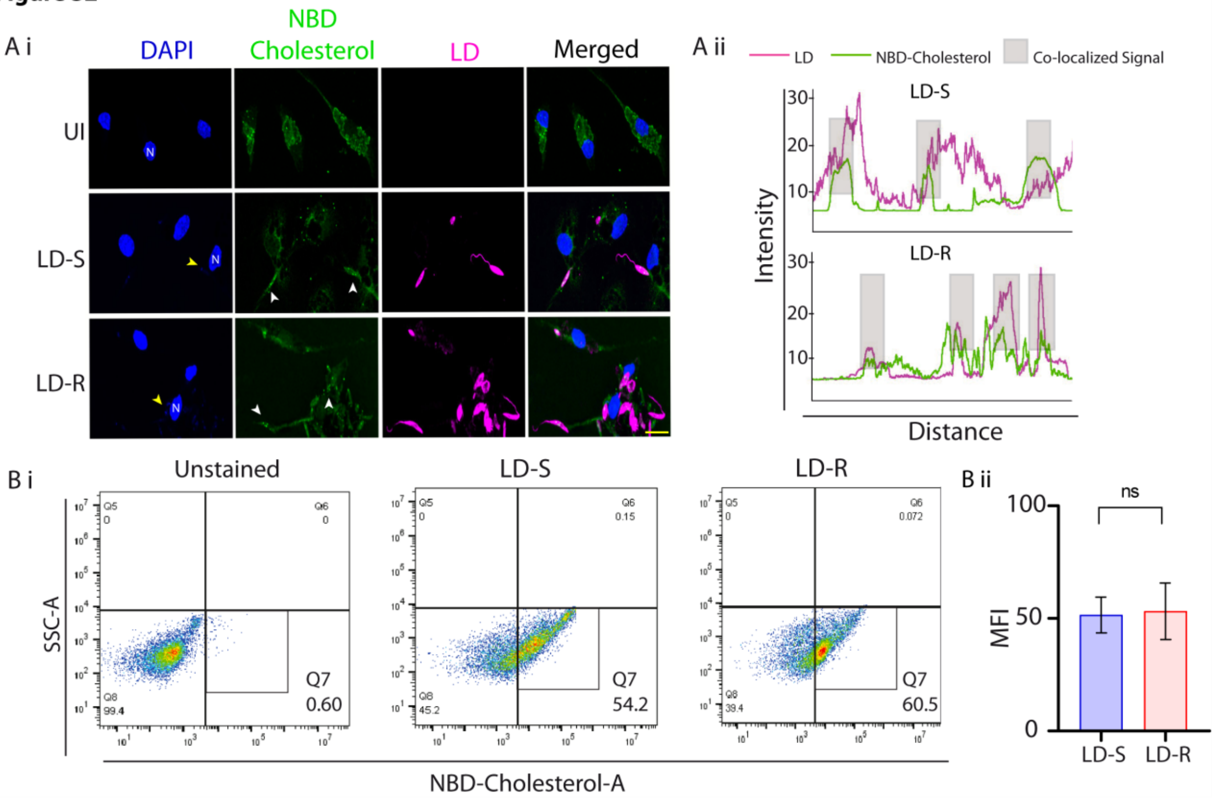

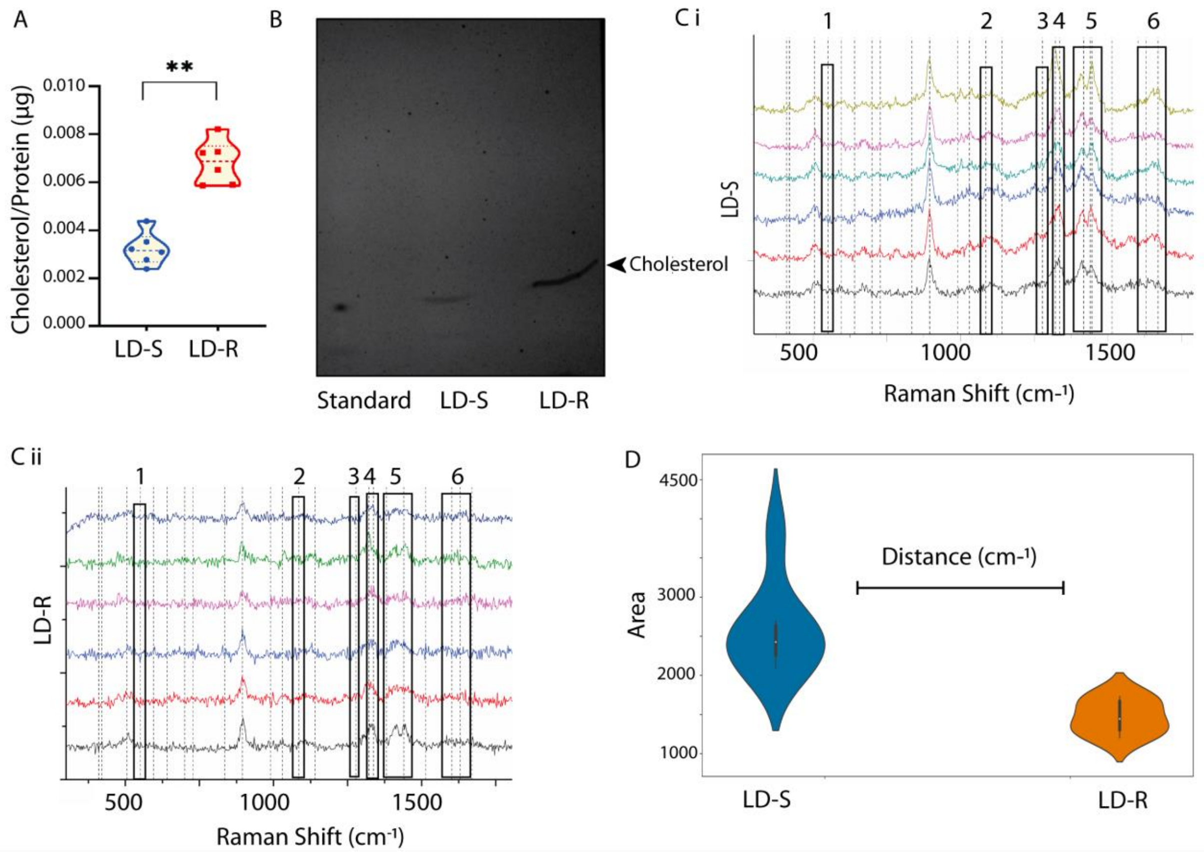

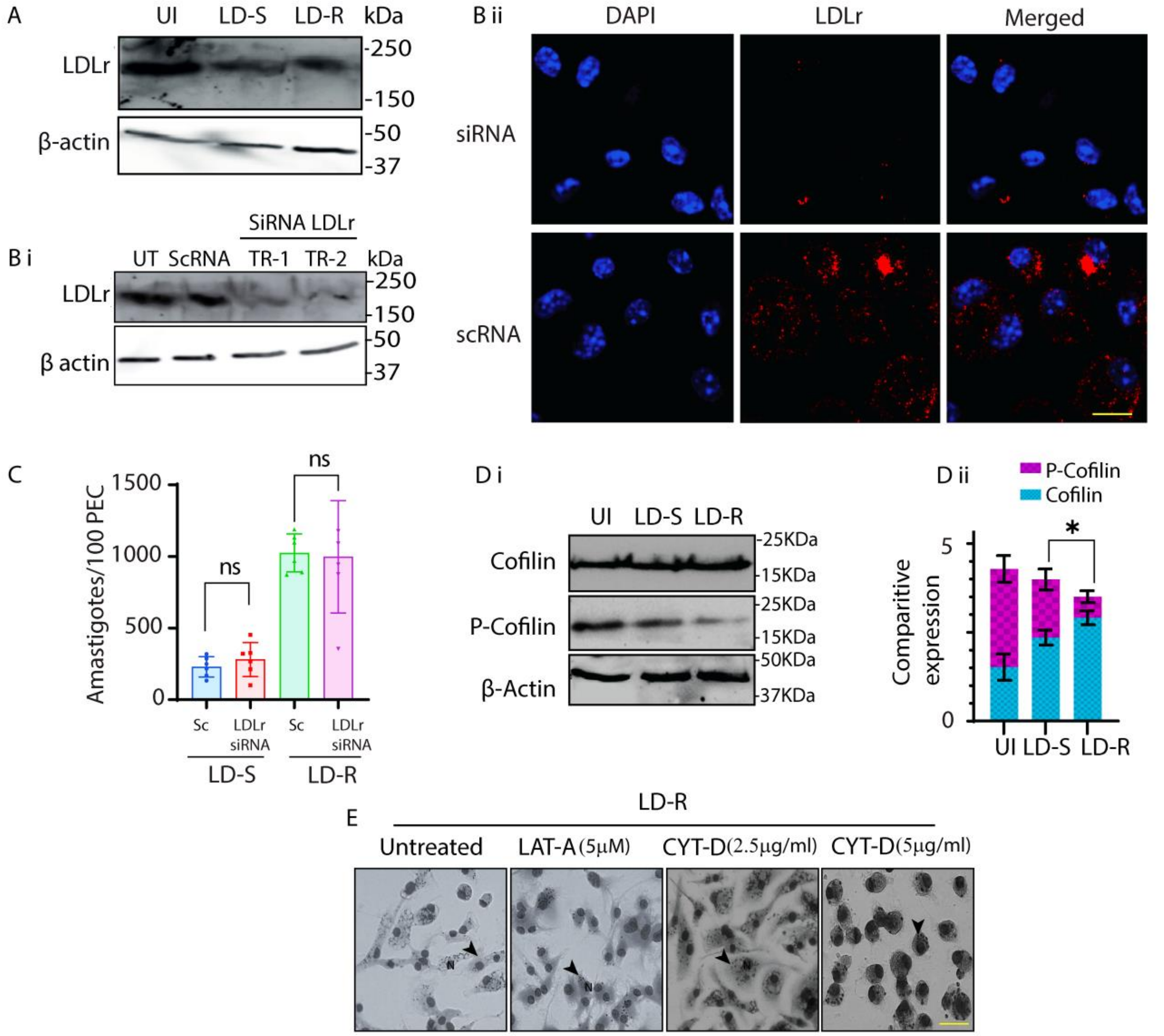

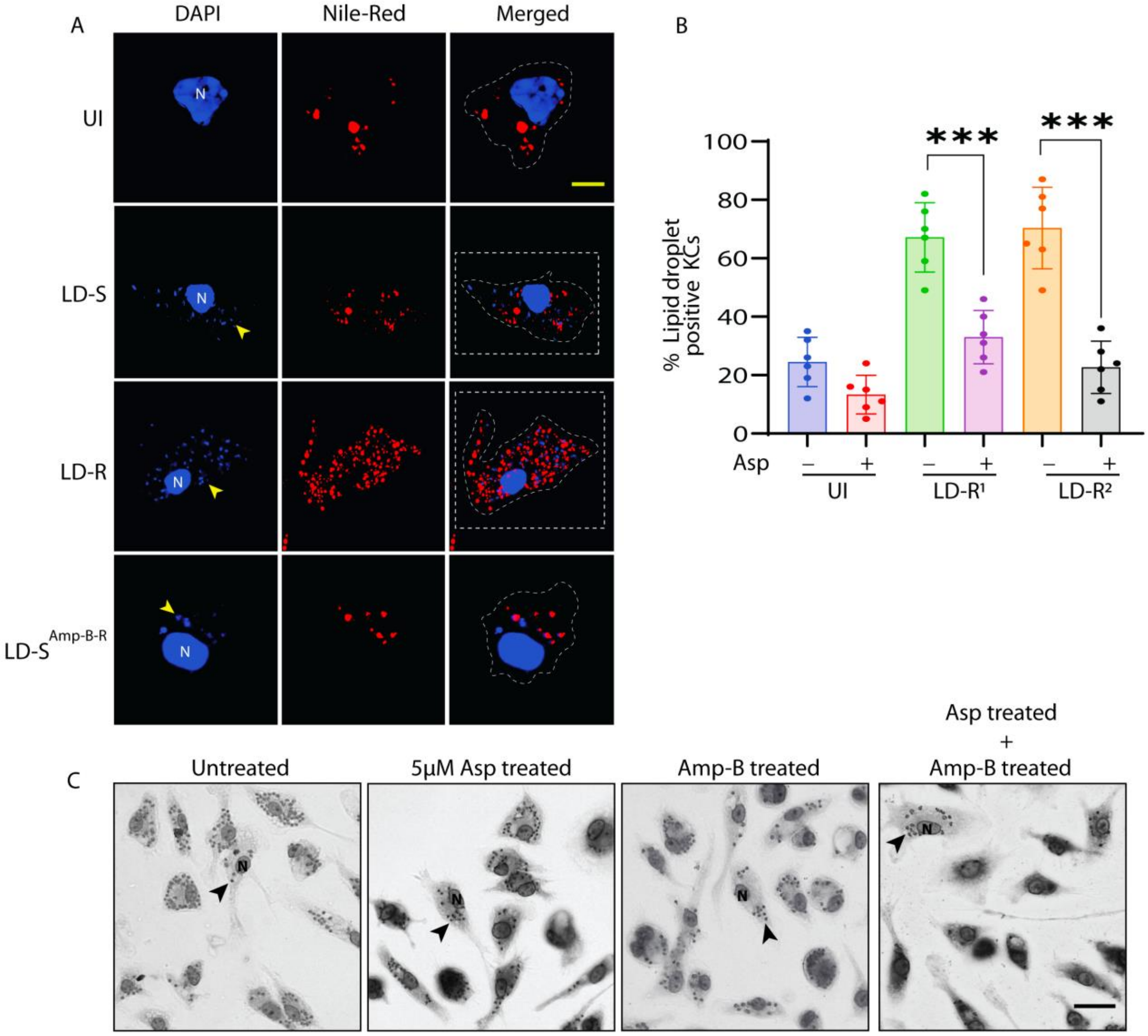
